# Emergent, cost-free surplus of core biosynthesis governs bacterial fitness

**DOI:** 10.64898/2026.07.17.739201

**Authors:** Huijing Wang, Dotan Goberman, Christopher Aldrich, Rami Pugatch, Fangwei Si

## Abstract

Cells often express essential components in excess of the levels required to sustain steady-state growth. The concept of protein reserve or surplus has recently emerged as a strategy for long-term cell fitness in bacteria, typically assumed to reduce steady-state growth as a trade-off for faster adaptation to a new environment. However, the origin and physiological consequences of such surplus remain unclear, particularly for core cellular processes including transcription, translation, and central metabolism.

Here, we establish a unified operational definition of surplus and quantify it for selected key components of transcription, translation, and central metabolism machinery in *Escherichia coli* at the single-cell level. By combining microfluidics, quantitative fluorescence imaging, and CRISPR interference-based gene knockdown, we demonstrate that substantial fractions of core biosynthetic components can be removed without affecting steady-state growth under nutrient-limited conditions. Unexpectedly, this surplus imposes no cost on steady-state growth, in contrast to prevailing trade-off models. Instead, we show that surplus emerges as a passive consequence of substrate limitation using a simple theoretical model, which is based on the universal autocatalytic-network structure of bacterial cells. Perturbation experiments confirm model predictions that surplus of transcription and translation machinery accelerates growth adaptation to nutrient-rich conditions, without affecting steady-state growth. Moreover, under slow-growth conditions, surplus suppresses cell death by reducing the risk of stochastic collapse of biosynthetic cycles, also in accord with our theoretical model prediction.

Altogether, our results identify surplus as an intrinsic property of core biosynthesis and a key determinant of long-term bacterial fitness in fluctuating environments.

## Introduction

A central goal in microbial physiology is to understand how bacterial cells balance rapid growth, adaptability, and survival in fluctuating environments [REFs: Schaechter 1958, Kjeldgaard 1958, Neidhardt 1960, Maaløe 1979, Bremer 1996, Scott 2010, Jun 2018, Mori 2017, Bruggeman 2023]. Among these processes, the control of bacterial growth under steady-state conditions has been extensively studied, for example, through the widely accepted resource allocation model [REFs: Scott 2010, Scott 2022]. This framework describes how cells allocate resources between the production of proteins and the synthesis of their building blocks under general constraints, such as translation capacity and macromolecular density [REFs: Scott 2022, Belliveau 2021, Chure 2025, Erickson 2016, Chure 2023, Kratz 2023, Droghetti 2025].

Building on the resource allocation model, a series of trade-off principles between steady-state growth and other physiological behaviors have been proposed [REFs: Mori 2017, Basan 2020, Balakrishnan 2021, Wu 2023, Bruggeman 2023, Choi 2026]. Across these studies, the concept of a protein “reserve”, or surplus, has emerged, whereby bacterial cells maintain excess amounts of specific proteins to facilitate processes such as growth adaptation. In this view, surplus is interpreted as a resource-consuming investment that improves certain physiological functions at the expense of the steady-state growth rate. For example, ribosomes and metabolic enzymes have been suggested to contain reserves that facilitate growth adaptation to new environments [REFs: Koch 1971, Mori 2017, Basan 2020, Balakrishnan 2021, Grigaitis 2022, Wu 2023, Pavlou 2025]. In addition to adaptation, overabundance of some essential low-copy proteins has been shown to enhance growth robustness by buffering fluctuations in gene expression [REFs: Lo 2024, Choi 2026, Goberman 2026]. Moreover, redundancy in genes and metabolic pathways has been studied as a strategy for buffering against genetic perturbations [REFs: Stelling 2004, Kitano 2004].

Despite increasing interest, the existence and physiological role of surplus in core biosynthetic processes remain unclear. Core biosynthetic processes, including transcription, translation, and central carbon metabolism, govern the majority of biomass production and are universally essential in all conditions [Fig. 1A]. Therefore, understanding whether these core biosynthetic processes have surplus and whether such surplus incurs costs in cellular resources is pivotal to understanding the coordination of growth, adaptation, and survival.

**Figure 1.**
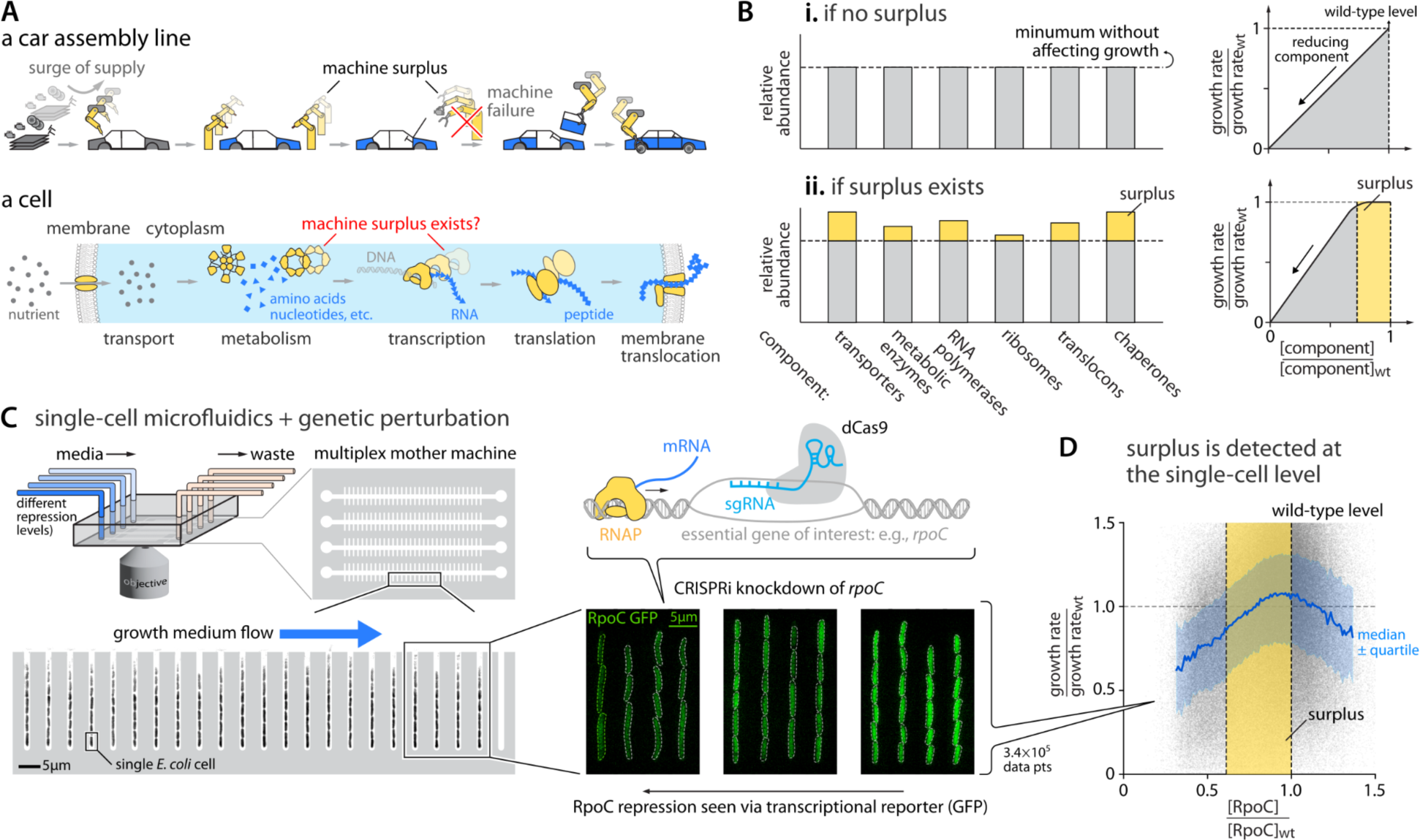
Surplus of core cellular components exists in single *E. coli* cells. **A.** Core biosynthetic processes can be viewed as part of an autocatalytic network [REFs: Roy 2021, Goberman 2026]. Each step may possess a surplus of substrates and/or catalytic machines, enabling rapid production upon increased substrate availability or buffering against failure of biosynthetic components [REFs: Koyuncuoğlu 2001, Sagron 2001]. **B.** Schematic illustrating the operational definition of surplus. **i.** In the absence of surplus for a given biosynthetic component (e.g., RNA polymerase), reducing its expression level (defined as copy number per cell volume) leads to an immediate decrease in steady-state growth rate. **ii.** In the presence of surplus, some reduction of the component’s expression level does not affect growth rate. **C.** We use a parallelized mother machine microfluidics platform in combination with CRISPR interference and fluorescent labeling to measure changes in growth rate while tuning the expression of essential components. Representative fluorescence images show gradual reduction of active RNA polymerase through CRISPR knockdown of its essential subunit ꞵ′, encoded by *rpoC*. **D.** The surplus of RpoC is quantified when plotting single-cell instantaneous growth rate as a function of RpoC expression level during steady-state growth, with both quantities normalized to the wild-type values. The yellow-shaded region indicates the surplus regime, corresponding to the length of the plateau region in the plot, defined as the expression level for which growth rate exceeds 90% of the wild-type level based on Hill-function fitting [Methods]. The calculation of surplus is robust across different methods for determining the plateau region [Figs. S2A and S2B, Methods]. Gray points represent individual measurements from single cells at a given time point, obtained using the mother machine platform (>340,000 data points). The blue line represents the median, with the interquartile range shown as shading. Some cells exhibit higher-than-wild-type RpoC expression and reduced growth, possibly reflecting proteome allocation constrained by extra RNAPs or other secondary effects, which are beyond the scope of this study.

Measuring surplus requires controlled perturbation of gene expression together with accurate measurement of growth. However, population-level reductions in growth rate may arise from both decreased biosynthetic capacity within individual cells and increased fractions of non-growing cells, which are difficult to distinguish using conventional batch-culture measurements [REFs: Hawkins 2020, Parker 2020, Donati 2021, Keren 2016, Otto 2024, McCain 2024]. Therefore, more precise characterization of surplus requires measurements that resolve distinct physiological states at the single-cell level.

Here, we address these challenges by combining single-cell measurements with controlled perturbations to establish a general and quantitative framework. We introduce an operational definition of surplus that enables direct experimental quantification and apply it to selected core components of central metabolism, transcription, and translation in *E. coli*. Our results reveal that the tested core biosynthetic components maintain an unexpectedly large surplus, particularly under nutrient-poor conditions. We find that surplus does not impose a measurable cost on the steady-state growth rate, in contrast to prevailing trade-off models, challenging the view that surplus represents a costly adaptive reserve. We further demonstrate that surplus of transcription and translation machinery accelerates growth adaptation to nutrient-rich conditions, and confers another key benefit – enhancing physiological robustness by suppressing cell death under slow-growth conditions. Our experimental observations are qualitatively explained by a simple model that is based on an autocatalytic network of biosynthesis. Together, our findings identify surplus as an intrinsic property of core biosynthesis and a key determinant of long-term bacterial fitness in fluctuating environments.

## Results

### 1. Surplus of core biosynthetic components exists in single *E. coli* cells

We operationally define the surplus of a core biosynthetic component as the maximal fraction of its copy number in the cell that can be removed without affecting the steady-state growth rate. Take, for example, RNA polymerase (RNAP) – a key component of the transcription machinery. As discovered by Nomura et al. [REF: Nomura 1986], 50% of RNAP in E. coli can be removed without altering steady-state growth rate, so half of the RNA polymerases in the cell are surplus. Operationally, to measure surplus of a cellular component, one can reduce its expression level, defined as the copy number per cell volume, and observe how much reduction will cause for the first time a subsequent reduction in the growth rate [Fig. 1B ii.]. If an essential molecular machine lacks surplus, even a small reduction in its expression level causes a measurable decrease in the growth rate [Fig. 1B i.].

To quantify surplus, we examined core biosynthetic components in *E. coli*, focusing on those involved in central metabolism, transcription, and translation. To this end, we characterized the relationship between two quantities: the reduced expression level of an essential biosynthetic component and the corresponding steady-state growth rate of the cell [Fig. 1B].

To measure the expression level of cellular components and growth rate across multiple conditions simultaneously, we use a parallelized version of the mother machine microfluidics platform [Fig. 1C, Fig. S1A] [REFs: Wang 2010, Thiermann 2024, Boesen 2024]. The mother machine enables continuous measurements of both single-cell growth rates and the expression levels of fluorescently labeled gene products within the same cells over time, thereby allowing reconstruction of growth-expression relationships across thousands of individual cells during an experiment. To systematically reduce the expression level of a target gene product, we employed a CRISPR interference system (CRISPRi) with validated specificity and tunability [Fig. 1C, Figs. S1F and S1G, Methods] [REF: Li 2016]. A fast-maturing green fluorescent protein (msfGFP) was inserted immediately downstream of the gene of interest, either as a transcriptional reporter or as a translational fusion, enabling quantitative measurement of gene expression using fluorescence imaging [Table S2, Methods].

We first examined the surplus of the transcription machinery, RNA polymerase (RNAP), which has previously been shown to contain a substantial inactive fraction [REFs: Nomura 1986, Cole 1987]. To do so, we grew E. coli cells in the mother machine under steady-state conditions while using CRISPRi to continuously repress *rpoB* or *rpoC*, both encoding essential components of RNA polymerase, across a range of expression levels [Figs. S2 and S3, Methods, SI figures].

The single-cell growth rate was quantified as the instantaneous rate of cell elongation, which is equivalent to the volume expansion rate in our experiment [REF: Kar 2025]. Gene expression level (copy number of gene product per cell volume) was measured as fluorescence intensity per cell volume (fluorescence concentration), with both obtained from the same cell at each time point [Methods]. By correlating growth rate and gene expression level across thousands of individual cells and multiple gene-reduction levels, we reconstructed the growth-expression relationship and thereby quantified surplus [Fig. 1D, Fig. S2, Methods]. Both growth rate and gene expression level were normalized to their wild-type values. Consistent with our definition, surplus was quantified as the length of the plateau region of growth rate as a function of the expression level, where growth remains unaffected despite reduced expression [Figs. 1B and 1D].

Under intermediate-growth conditions (minimal glucose medium doubling time ≈ 60-80 minutes), both RpoB and RpoC exhibit a similar and substantial surplus [Fig. 2A, Figs. S2 and S3]. The similarity between RpoB’s and RpoC’s surplus is consistent with the fact that both their genes are on the same operon and perturbation of either gene similarly affects the total RNAP copy number. Overall, the single-cell growth rate remains essentially unchanged even when RpoB or RpoC is reduced to approximately 60% of the wild-type level, indicating that nearly half of the total copies of RNAP are not required to sustain steady-state growth.

**Figure 2.**
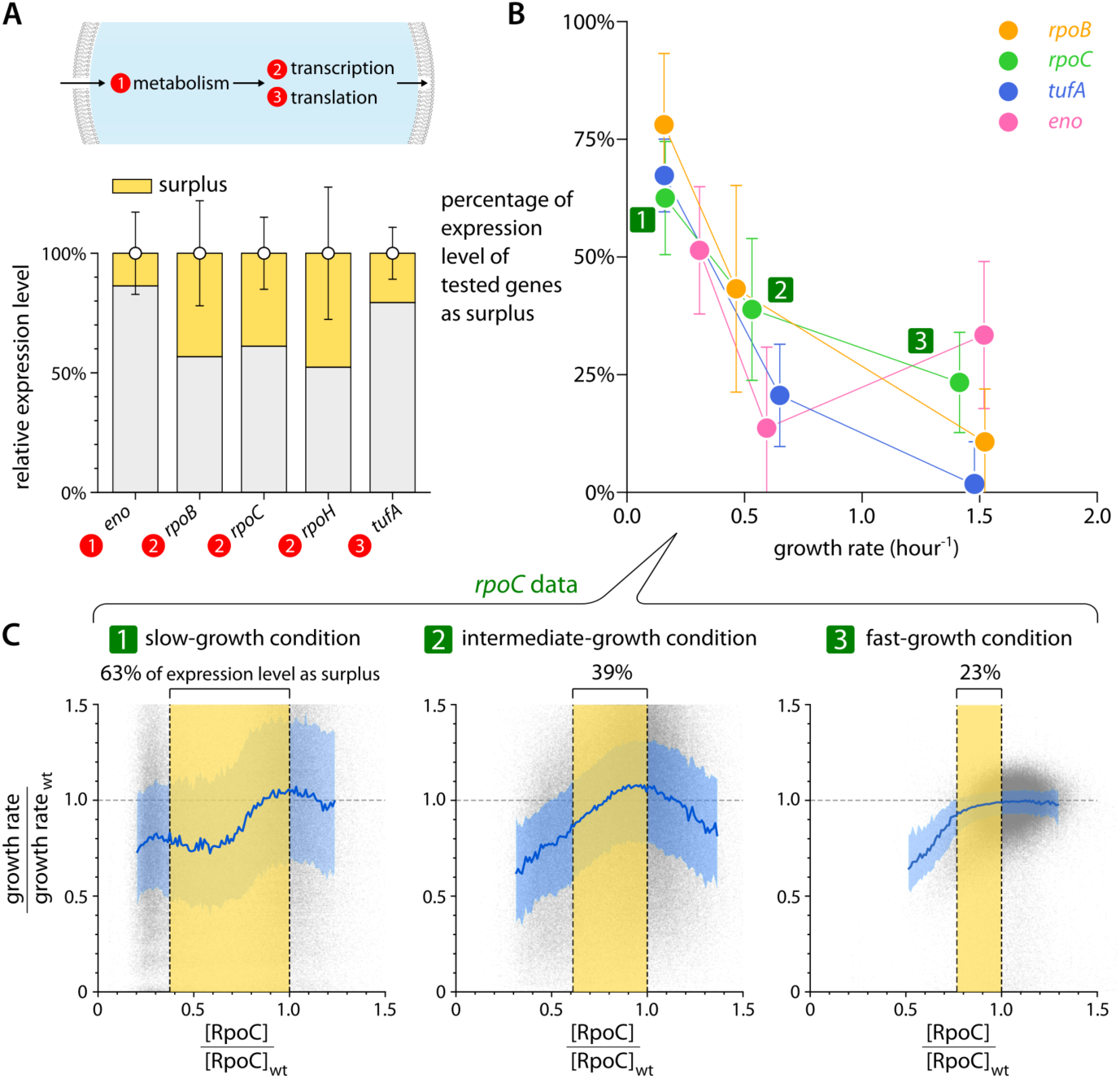
Surplus of different core biosynthetic components systematically increases under slow-growth conditions. **A.** All tested core biosynthetic components possess a surplus under an intermediate-growth condition. Yellow shading represents the surplus, defined as a fraction of the wild-type expression level of each gene. Error bars represent the standard deviation in wild-type expression level. The stress-associated sigma factor RpoH exhibits a larger surplus (∼50%), consistent with its limited requirement under normal growth conditions [also see Figs. S2, SI figures]. **B.** The magnitude of surplus in various core biosynthetic components systematically increases as the growth rate decreases under nutrient-poorer conditions. RpoB and RpoC, both essential subunits in RNAP, exhibit similar levels of surplus. Error bars are defined as in **A. C.** Using RNAP components as an example, the growth-expression relationship for RpoC shows a larger plateau region in slow-growth conditions, corresponding to surplus. In the slow-growth condition (MOPS acetate), surplus reaches 63% of the wild-type expression level, compared to 39% and 23% in intermediate (MOPS glucose) and fast-growth (MOPS rich glucose) conditions, respectively. Data are presented as in Fig. 1D. See Figs. S2 for additional genes.

### 2. Different core biosynthetic components all exhibit significant surplus

Having quantified a substantial surplus in the transcription machinery, we next asked whether this phenomenon is specific to RNA polymerase or instead reflects a general feature of core biosynthetic processes. To determine whether surplus is a general property of core biosynthetic processes, we measured the surplus of representative components involved in translation and central metabolism [Fig. 2A]. For translation, we perturbed the essential elongation factor EF-Tu, encoded by *tufA* in a standalone operon, owing to its genetic accessibility [REF: Karp 2023]. Perturbing TufA reduces translational capacity, thus effectively altering the copy number of functional ribosomes. For central metabolism, we examined enolase, encoded by *eno*, the sole essential enzyme catalyzing the conversion of 2-phosphoglycerate to phosphoenolpyruvate in glycolysis [REF: Karp 2023].

Under the same intermediate-growth condition (minimal glucose medium) as for the RNAP experiment, all tested components exhibited measurable surplus, with approximately 10-40% of the total copy number can be removed without affecting steady-state growth [Fig. 2A, Figs. S2 and S3, Methods, SI figures]. Notably, we did not observe a clear correlation between the magnitude of surplus and either the function or the expression level of the component, suggesting that surplus is a widespread property of core biosynthetic processes rather than merely determined by protein class or expression level.

### 3. Surplus of core biosynthetic components increases under slow-growth conditions

Having established that surplus is broadly present across biosynthetic systems, we next asked whether surplus itself depends on growth conditions. The composition of biosynthetic components is known to vary systematically with growth rate, which itself depends on nutrient quality [REF: Bremer 1996, Scott 2022], raising the question of whether surplus varies across different nutrient conditions. To investigate this, we quantified surplus across a range of nutrient conditions spanning doubling times from 27 minutes to more than 3 hours [Fig. S2, Table S3, Methods, SI figures].

Notably, the magnitude of surplus increases systematically at slower growth rates under nutrient-limited conditions. For example, the surplus of RpoC is approximately 23% of the wild-type expression level in fast-growth conditions (MOPS rich glucose), increases to 39% in intermediate conditions (MOPS glucose), and further expands to 63% in the slow-growth conditions (MOPS acetate) [Figs. 2B and 2C].

The trend of a larger surplus under slower-growth conditions is not limited to the transcription machinery. All other tested components – including those involved in translation and central metabolism – exhibit increasing surplus under slower-growth conditions, reaching 50-80% of their wild-type levels under the slowest-growth conditions examined [Fig. 2B]. The dependence of surplus on growth condition is robust across different methods used to quantify surplus [Fig. S4], indicating that the observed trend reflects an intrinsic and general property of cellular physiology. Together, these results reveal that surplus is not only widespread but systematically amplified under nutrient limitation.

### 4. Surplus arises as a passive consequence of substrate limitation

The observation that such a large surplus imposes no cost on steady-state growth challenges the prevailing view that surplus reflects a resource allocation trade-off. If surplus were costly, reducing it would be expected to increase growth rate, which we do not observe. This apparent discrepancy raises a fundamental question: if surplus does not incur a cost, what determines its magnitude and origin?

The systematic increase of surplus under nutrient limitation suggests that surplus may arise from the same constraints that limit growth. To explain the biological origin of surplus, we developed a quantitative framework based on the autocatalytic-network structure of cellular biosynthesis [see details in SI notes A and B] [REF: Roy 2021]. In this framework, core biosynthetic processes are organized as interconnected autocatalytic cycles in which the transcription-translation machinery synthesizes all proteins by consuming substrates. Importantly, the transcription-translation machinery also replicates copies of itself. A basic feature of this model is that, for any catalyst-substrate reaction, the production rate is limited by whichever component is in shorter supply – either the catalyst (enzyme) or the substrate [SI notes A and B] [REF: Roy 2021]. This formulation differs from many existing models, which typically assume that biosynthetic rates depend primarily on enzyme copy number.

The central idea of this model is that biosynthetic reactions can be limited by either substrate copy number or enzyme copy number, depending on growth conditions. To demonstrate how this model works, consider the transcription process as an example [REF:Roy 2021]. The dynamics can be written as,

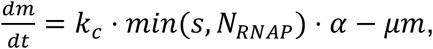

where the rate constant *k*_*c*_ sets the timescale of transcription, *m* is the number of mRNA molecules of a given gene, *s* is the copy number of substrate (nucleotides), *N*_*RNAP*_ is the number of RNA polymerases, *α* is the fraction of RNAP allocated to that gene, and *μ* is the mRNA degradation rate. In nutrient-poor conditions (slow-growth), at a given moment, if the substrates are too scarce to feed all RNAPs, namely, *s* < *N*_*RNAP*_, then the number of actively transcribing RNAPs is no more than the number of free nucleotides [Fig. 3A i]. In this regime, mRNA production depends only on substrate copy number and is independent of RNAP copy number, such as, 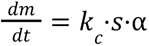. Conversely, in nutrient-rich conditions, where the substrates are abundant and saturating all RNAPs, *s* ≥ *N*_*RNAP*_, we have, 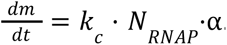. The relation is reduced to the case in which mRNA production depends on RNAP copy number, as is typically employed in many bacterial growth models [Fig. 3A i].

**Figure 3.**
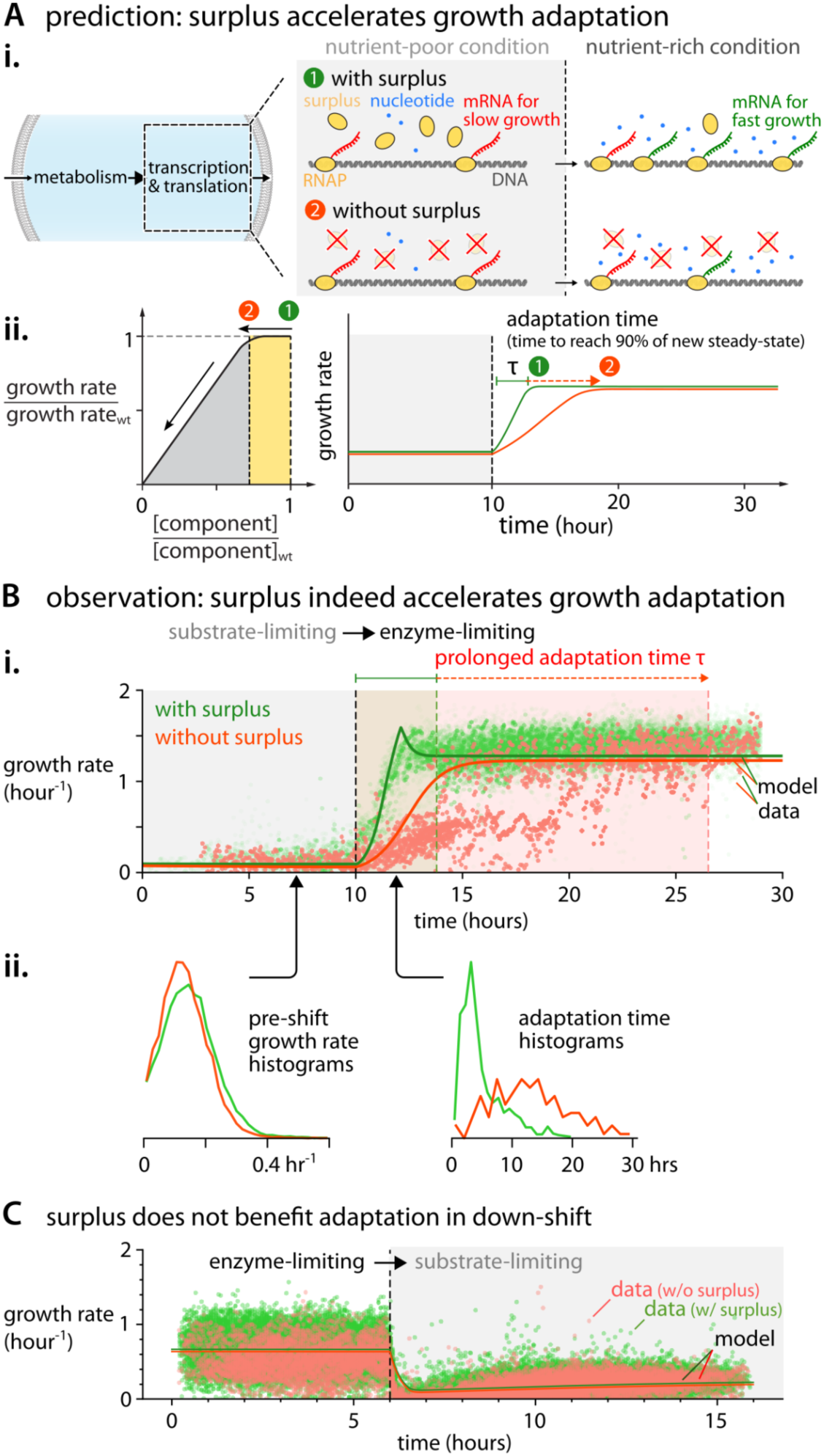
Surplus of transcription and translation machinery facilitates growth adaptation to nutrient-rich conditions. **A.** Model predictions for surplus’ benefit in growth adaptation. **i:** In the presence of surplus, transcriptional rate is initially limited by substrate availability (e.g., nucleotides), such that a fraction of RNA polymerase (RNAP) remains unused. Following a nutrient upshift, when substrates become abundant, this pre-existing surplus can be rapidly mobilized for global transcriptional remodeling, thus accelerating growth. In the absence of surplus, transcription becomes immediately limited by RNAP copy number, resulting in slow growth adaptation. Non-specifically DNA-bound RNAP is represented as free RNAP for simplicity. **ii:** Removing surplus is predicted to have little effect on steady-state growth rates, but to significantly increase the adaptation time, which is defined as the time taken to almost reach the new steady-state growth rate. **B.** The nutrient up-shift experiment is in reasonable agreement with the prediction of our simplified model. Time-course measurements show that reducing RpoC surplus (red dots in **ii.**) leads to a pronounced slowdown in growth adaptation following a shift from a slow-growth condition (MOPS acetate with arginine) to a fast-growth condition (MOPS rich glucose), compared to cells with unperturbed surplus (green dots in **i.**). A more detailed modeling approach, beyond the scope of this research, is required to capture the behaviors of different cell subpopulations that respond at different timescales to the upshift. Reducing surplus also leads to significant cell death, as studied in Section 6, resulting in fewer data points. The colored shaded regions indicate the adaptation time τ, defined as the time required to reach 90% of the maximum growth rate [Methods]. The pre-shift growth rate histograms remain unchanged with or without surplus, but the adaptation time histogram shifts to larger values when surplus is reduced (**ii.**), in contrast to trade-offs on growth rate by surplus. The single-cell adaptation time in the histogram is measured by tracing single-cell lineages throughout the nutrient shift. Data points in **i.** represent the instantaneous single-cell growth rate (> 22,000 measurements). Solid lines correspond to model calculations, with surplus as the only free parameter [SI notes C]. The model also captures the transient overshoot in growth rate following the shift. **C.** Reducing RNAP surplus does not significantly affect adaptation rate following a downshift from a nutrient-rich (MOPS glucose with 11 amino acids) to a nutrient-poor (MOPS acetate) condition, consistent with model predictions.

Within this framework, the surplus of RNAP is calculated as,

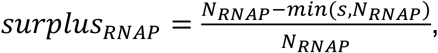

arising naturally as the fraction of enzyme molecules that are not engaged in catalysis due to insufficient substrate availability [see details in SI notes A and B]. Importantly, the surplus results from incomplete enzyme utilization rather than active overproduction of the enzyme, and therefore does not impose an additional cost on growth. The same reasoning applies to other core biosynthetic processes, including translation and central metabolism, in which substrates such as amino acids or metabolic intermediates can become limiting. Accordingly, surplus emerges as a passive consequence of substrate-limited kinetics in an autocatalytic network, providing a unified explanation for both the widespread occurrence of surplus and its systematic increase under nutrient limitation.

### 5. Surplus accelerates growth adaptation without affecting steady-state growth

An key prediction of our model is that the surplus accelerates the adaptation of cell growth to a new steady state upon a nutrient upshift. Using transcription as an example, in nutrient-poor conditions where substrates (nucleotides) are limiting, RNAP surplus does not contribute to steady-state growth. However, upon a shift to nutrient-rich conditions, many genes required for fast growth recruit new RNAPs, and substrates are sufficiently supplied to saturate RNAPs. After the nutrient upshift, RNAP surplus can be immediately mobilized to transcribe fast growth-related genes, including ribosomal proteins and RNAP’s genes, thereby increasing the growth rate and accelerating the adaptation to the new steady state [Fig. 3A ii., SI notes C]. In the absence of RNAP surplus, additional RNAP must first be synthesized before transcriptional capacity can increase, thereby slowing down adaptation.

We used the autocatalytic-network model described above to theoretically explain how surplus shortens the adaptation time upon shifting to a nutrient-rich environment. To extend the model beyond steady-state growth, we incorporated a dynamical control law that modifies the allocation of the transcription-translation machinery towards its own reproduction when the external limitation of nutrients is suddenly lifted. The controller attempts to lock the allocation parameter to a fixed ratio between the supply of available nutrients, e.g., amino acids, which suddenly becomes higher, and its demand by the autocatalytic cycle that replicates the transcription-translation machinery [SI notes A and B]. Our highly simplified model qualitatively shows that repressing surplus of transcription or translation machinery can prolong the adaptation period several-fold during a nutrient upshift, without affecting the steady-state growth rate before or after the shift [model lines in Fig. 3B i.], using the same parameters for the controller gain [SI notes C]. Due to the simplicity of the model, we can track the reason for the shortening of the adaptation time. Adaptation time shortens since the presence of surplus of core transcription-translation machinery brings the cellular composition closer to the new steady-growth cellular composition after the upshift, requiring less time to complete the missing molecular machines (e.g., RNA-polymerase, ribosomes, or other growth-required enzymes).

To test the prediction that surplus accelerates growth adaptation, we performed a nutrient upshift experiment by switching cells from a slow-growth condition (e.g., MOPS sodium acetate with arginine as the sole nitrogen source, doubling time ≈ 4.3 hours) to a fast-growth condition (e.g., MOPS glucose supplemented with synthetic rich nutrients including amino acids and nucleotides) [Fig. 3B, Figs. S1C-S1E, S5, S6, and SI figures for other choices of growth media, Table S4, Methods].

Consistent with the prediction that surplus contributes to faster adaptation, reducing the surplus of RpoC or RpoB – either throughout the shift experiment [Fig. 3B, Figs. S5 and S6, SI figures] or only during the pre-shift phase [SI figures] – significantly slows down growth adaptation [Fig. 3B]. In contrast, both the pre-shift and post-shift steady-state growth rates remain largely unchanged [Fig. 3B ii.], in agreement with the simple theoretical model [Fig. 3B i. solid lines, Figs. S5 and S6, SI figures]. The decoupling between steady-state growth rate and adaptation demonstrates that surplus is not governed by a trade-off with growth rate, but instead reflects the underlying limitation imposed by substrate availability. The transient overshoot in growth observed during adaptation is also captured by the model, consistent with regulatory feedback dynamics incorporated in the framework [SI notes C and D]. Reducing surplus caused a subset of cells to exhibit adaptation times substantially longer than predicted by the deterministic model, suggesting that surplus may also influence adaptation through mechanisms not captured by the current framework. Perturbing surplus also leads to significant cell death, as examined in Section 6, resulting in fewer growing cells following the shift [Fig. 3B i]. A similar slowdown in adaptation is observed for perturbation of the translation-related protein TufA, again supporting the generality of the model [SI figures].

Another prediction is that surplus in transcription machinery should not enhance adaptation during nutrient downshifts, where substrate limitation remains dominant. Consistent with this prediction, experimental measurements show that cells with reduced RNAP surplus do not exhibit prolonged adaptation following a downshift [Fig. 3C solid lines, Figs. S9, SI figures].

Finally, we predict that surplus in central metabolic enzymes should have little effect on growth adaptation, as these enzymes are not strongly limiting under either slow- or fast-growth conditions [Fig. S7A]. Consistent with this prediction, perturbation of the essential metabolic enzyme enolase does not significantly affect adaptation dynamics [Fig. S7 and S8, SI figures]. Importantly, despite their distinct functional roles in adaptation, the surplus of all tested components does not influence steady-state growth rate. Taken together, the experimental results and quantitative model demonstrate that surplus in transcription and translation accelerates adaptation during nutrient upshifts without imposing a cost on steady-state growth, in contrast to previously reported growth-adaptation trade-offs [REFs: Mori 2017, Basan 2020, Balakrishnan 2021, Wu 2023, Bruggeman 2023, Choi 2026].

### 6. Surplus reduces cell death by suppressing stochastic collapse of biosynthetic autocatalytic cycles

While enhanced adaptation speed provides a clear fitness advantage following environmental shifts, the widespread presence of surplus under slow-growth conditions suggests an additional important physiological function beyond adaptation. Therefore, we asked whether surplus also contributes to physiological robustness by buffering stochastic fluctuations in core biosynthetic processes [REF: Goberman 2026].

As mentioned, RNAP in *E. coli* at slow growth rate has 50% or more surplus, i.e., at least 50% RNAP can be removed without altering steady-state growth rate. However, reducing the number of RNAP at slow growth rates comes with a price — an increase in the probability of cell death, as we now explain.

For example, although the total number of RNA polymerases (RNAP) per cell is typically a few thousand under intermediate growth conditions, only a handful are engaged in transcribing a given gene at any moment, including genes encoding RNAP’s own subunits such as *rpoB* or *rpoC*.

Removing RNAP surplus, which is equivalent to removing 50% or more of the total copy number, amplifies relative fluctuations in the expression of its own genes. In turn, these fluctuations increase the probability of disruption of RNAP autocatalytic production, increasing the probability of collapsing its autocatalytic cycles which can ultimately lead to cell death [see model details in SI notes E] [REF: Goberman 2026] [Fig. 4A].

**Figure 4.**
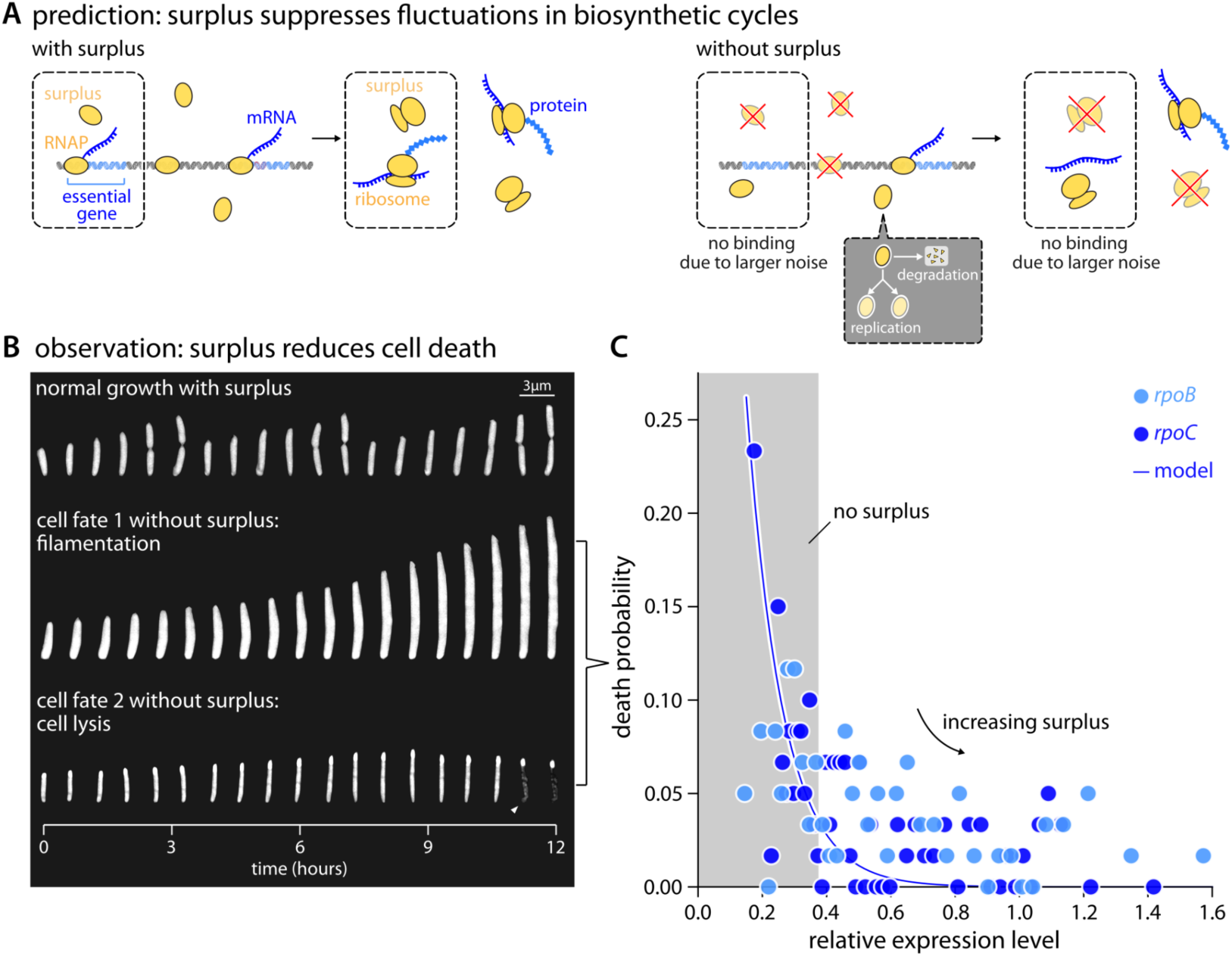
Surplus of core biosynthetic components reduces cell death by suppressing stochastic collapse of essential biosynthetic cycles. **A.** According to our model, although the total copy number of RNAP or ribosomes is high, only a small number of these components are typically engaged in the production of individual essential gene products at any given time. Reducing surplus further decreases these numbers, amplifying stochastic fluctuations in biosynthetic cycles and increasing the likelihood of collapse. The collapse is more likely under slow-growth conditions, where replication and degradation rates of these components become comparable [REF: Goberman 2026]. **B.** With reduced RNAP surplus under steady-state conditions, cells exhibit higher frequencies of filamentation or cell lysis, with filamentous cells eventually dying. Phase-contrast cell images are snapshots from 12-hour time-lapse experiments; the triangle marks the moment of cell lysis. **C.** The death probability under steady-state conditions, defined as the combined probability of cell lysis and filamentation, increases as the expression level of RpoB or RpoC is reduced. Model predictions (solid lines) quantitatively capture the observed increase in death probability. The only fitted parameter, the RNAP fraction allocated to the *rpoBC* operon, agrees well with literature values [see model details in the SI notes E]. The vertical dashed line denotes the expression level corresponding to the boundary of surplus; to the right of this boundary, surplus reduces the probability of cell death.

To test the role of surplus in suppressing cell death, we measured cell survival under conditions in which RNAP surplus was perturbed while steady-state growth was maintained. We find that reducing RNAP surplus significantly increases the probability of cells that undergo physiological failure, even though the growth rate of surviving cells remains largely unchanged. The physiological failures manifest as cell lysis or cell filamentation, the latter eventually leading to cell death [Fig. 4B]. Quantitatively, the increase in cell death probability – defined as the combined probability of lysis and filamentation – upon perturbation of RpoC or RpoB is well captured by the model, using parameters consistent with independent measurements [Fig. 4C, SI notes E].

The role of surplus in suppressing fluctuations further predicts that the probability of cell death should be higher under conditions where fluctuations in the copy numbers of core components are enhanced. Consistent with this prediction, we observe elevated cell death probabilities during nutrient upshifts and during recovery from carbon starvation [Figs. 4C and S10A], where the probability of RNAP depletion could be augmented via stronger competition for binding to different sigma factors upon environmental shifts [REFs: Klumpp 2008, Yan 2024]. Similar enhanced cell death probabilities are observed for perturbations of the ribosomal component TufA, and the metabolic enzyme enolase [Figs. S10B and S10C], indicating that the effect of surplus on physiological robustness extends beyond transcription.

Together with the adaptation results, the robustness results indicate that surplus contributes to two independent determinants of long-term fitness: rapid adaptation and resistance to stochastic physiological failure. Importantly, both benefits arise without affecting steady-state growth rate, further supporting the conclusion that surplus is not associated with a growth cost but instead reflects an intrinsic feature of biosynthetic network organization.

## Discussion

Here, we establish an operational definition of surplus that is experimentally tractable and enables systematic comparison across cellular components and growth conditions. Our measurements provide a quantitative framework for incorporating surplus into physiological models of cellular growth and adaptation. Previous frameworks typically assume that biosynthetic flux is determined jointly by enzyme abundance and substrate availability, e.g., in models of ribosome-mediated protein synthesis or metabolic flux analysis [REFs: Scott 2010, Erickson 2016, Lin 2018, Chure 2023, Kratz 2023, Droghetti 2025, Orth 2010a, Orth 2010b, Monk 2013, Dourado 2023, Bruggeman 2023]. In contrast, our results show that core biosynthetic components exhibit substantial surplus, particularly under nutrient-limited conditions. Incorporating surplus into physiological models may improve predictions of cellular behaviors while providing a more realistic description of biosynthetic limitations under nutrient-poor conditions.

Our finding that large surplus does not impose a measurable cost on steady-state growth contrasts with widely proposed trade-offs between growth and adaptation [REFs: Mori 2017, Basan 2020, Balakrishnan 2021, Wu 2023, Bruggeman 2023, Choi 2026]. This apparent inconsistency is resolved by our model, which explains surplus as a passive consequence of incomplete enzyme utilization rather than active overproduction. As a result, reducing surplus does not increase growth rate, because growth remains limited by substrate availability.

The prevalence and magnitude of surplus across core biosynthetic processes suggest additional physical constraints shape cellular physiology under nutrient limitation. These constraints may include limits on enzymatic activity, nutrient uptake and transport [REFs: Koch 1982, Koch 1996, Ferenci 1996], macromolecular crowding in cytoplasm and cell membranes [REFs: Zhuang 2011, Szenk 2017], and geometric constraints such as surface-to-volume ratio of the cell [REF: Chure 2025, Trickovic 2026]. Therefore, surplus can be understood as an emergent property of biosynthetic systems operating under multiple constraints, rather than as a specifically regulated reserve. Further work will be needed to disentangle the relative contributions of these physical limitations.

Beyond its origin, surplus has important functional consequences for cellular fitness. Our results show that the surplus of transcription and translation machinery facilitates rapid growth adaptation following nutrient upshifts, while also reducing the likelihood of cell death by buffering stochastic fluctuations in biosynthetic processes [Figs. 3 and 4]. Moreover, we observed the role of the essential metabolic enzyme enolase’s surplus in maintaining morphological robustness and survivability upon environmental changes. Enolase catalyzes the conversion of phosphoenolpyruvate (PEP), a key precursor for peptidoglycan synthesis, which could explain its role in maintaining cell envelope integrity and survival, with the mechanistic link remaining to be investigated. These findings suggest that surplus simultaneously contributes to both adaptability and robustness, two key determinants of fitness in fluctuating environments.

From an evolutionary perspective, the magnitude of surplus may reflect the balance between these fitness benefits and the underlying constraints that give rise to it. For example, enhanced adaptation during nutrient upshifts could be advantageous in environments characterized by feast-and-famine cycles, such as the mammalian gut [REFs: Koch 1971, Mori 2017]. Similarly, the role of surplus in suppressing cell death in nutrient limitation may confer advantages during prolonged environmental stress. Therefore, surplus may not be an actively optimized strategy, but rather an intrinsic property of biosynthetic systems that is subsequently shaped by physical constraints under evolutionary pressures.

Overall, our results identify surplus as a general and emergent property of the core biosynthetic network that arises from substrate-limited kinetics and enhances both adaptation and robustness without imposing a measurable cost on steady-state growth. Rather than representing a costly adaptive reserve, surplus emerges naturally from the operation of biosynthetic autocatalytic networks under nutrient limitations. Our findings on surplus provide a unified framework linking molecular constraints, physiological robustness, and long-term bacterial fitness in fluctuating environments.

