## Supplementary material for "Emergent, cost-free surplus of core biosynthesis governs bacterial fitness": Methods, Supplemental figures and notes

#### Materials and methods

##### 1. Bacterial strain construction

Strain background

The *Escherichia coli* (*E. coli*) strains used in the mother machine experiments are derived from the K-12 MG1655 background. The strain is a low-motility *E. coli* MG1655 strain (Lyons et al., 2011) that was previously fully sequenced and suitable for the mother machine microfluidic experiments (Si et al., 2017, 2019).

Construction of the CRISPRi strains

In this study, we used a tunable CRISPR-Cas interference system, CRISPRi, for precise and continuous titration of target gene expression (Li et al., 2016). The CRISPRi system employs a tunable arabinose operon promoter $P_{BAD}$ to quantitatively control the expression of a nuclease-deficient Cas9 (dCas9), which binds to the target gene with the guidance of a small guide RNA (sgRNA) under constitutive expression, thereby interfering with transcription. Higher arabinose levels in the media enhance dCas9 expression, leading to stronger inhibition of targeted gene expression (Li et al., 2016). The expression of dCas9 alone, in the absence of any single-guide RNA (sgRNA), does not have a measurable impact on cell growth.

The background strain expresses dCas9 from $P_{BAD}$ and harbors DNA recombineering pSIM18 plasmids, with the arabinose transporters *araE* and the lactose inhibitor gene *lacI* deleted. It also has a counter-selectable marker, tetA-sacB, under a constitutive promoter, which is designed to be replaced by the specific sgRNA (Li et al., 2016). Constructing a parent strain is a one-step recombineering process, where a co-transcriptional reporter gene, *msfGFP,* is inserted at the end of the target gene on the cell chromosome. In the parent strain, the target gene expression is not regulated by arabinose concentration in the media. The successful insertion of the reporter gene was confirmed via both agar pad fluorescence imaging and Sanger sequencing before use for the mother machine experiments. A CRISPRi strain is generated from the parent cells by replacing the tetA-sacB cassette with an sgRNA fragment targeting essential genes via another step of recombineering. The sgRNA insertion and gene expression inhibition due to the inducible CRISPRi system were confirmed by agar pad fluorescence imaging before the mother machine microfluidic experiments. A list of constructed cells is available in Table S2.

Construction of the strain for gene overproduction

To construct a plasmid with inducible *rpoD* expression, *rpoD*, the transcriptional reporter mCherry, and the plasmid backbone with an inducible promoter $P_{LAC}$ were assembled via Gibson assembly reaction. The inducible *rpoD* plasmid was then transformed into *rpoC* tCRISPRi cells (Table S2). The plasmid construction and *rpoD* inducibility were confirmed with both plasmid sequencing and agar pad fluorescence imaging prior to mother machine experiments.

#### 2. Procedures of mother machine experiments

Fabrication and preprocessing of microfluidic devices

The mother machine enables precise control over cellular microenvironments, allowing either steady-state growth or controllable shifts between conditions. Each main trench on the mother machine microfluidics contains 4000 individual channels. It also enables quantitative measurements—such as growth rate and fluorescence-based expression levels—from the same cells, and supports the reconstruction of growth-expression relationships for thousands of individual cells within a single experiment. (Wang et al., 2010, Thiermann et al., 2024, Boesen et al., 2024). PDMS devices were fabricated as described in Wang et al (Wang et al., 2010). In steady-state growth experiments, for each main trench on a PDMS device, an inlet hole was used to allow fresh media flow in. In nutrient up-shift and downshift experiments, for each main trench, two inlet holes were used for media switching. Fig. S1 illustrates the configuration of the mother machine and syringe pump.

The fabricated PDMS devices were then bound to a WillCo-dish (WillCo Wells HBST-5040) glass bottom. Before use, machine devices were preprocessed in a plasma cleaner (Harrick Plasma PDC-32G) for 10 minutes at 0.8 Torr, baked at 80 ℃ for 10 minutes, and passivated with 0.5 mg/mL bovine serum albumin fraction V (BSA, Roche Diagnostics, CAS-No. 9048-46-8) for about 15 minutes. After preprocessing, the mother machines were ready to load cells.

Cell preparation

Cells constructed from the *E. coli* strain K-12 MG1655 background were used in this study (Table S2). Prior to each time-lapse imaging experiment, cells were picked from glycerol stock at -80 ℃ and spread onto an agar plate. After overnight incubation at 30 ℃, Cells were picked from agar plates and seeded into 1 mL LB media (MP Biomedicals, LLC, SKU: 113002122-CF). The agar plates were wrapped with parafilm and kept at 4 ℃. Plates older than 7 days were discarded. The cultures were then shaken for 12-18 hours at 30 ℃ in a water bath shaker. Based on the experimental conditions, we diluted the liquid cultures in the desired growth media until cell growth reached the exponential phase. 2 mL of the liquid culture was used to load one main trench of the multiplex mother machine device. The cultures were then concentrated 10- to 100-fold and injected into one mother machine's main trench with a 10 μl micropipette. With both inlets and outlets of the trenches sealed with small pieces of scotch tape, the cells were loaded into growth channels by centrifuging the mother machine device using a customized centrifuge, spun at 500 xg for 5 min. Fresh media were then infused with a syringe pump. See Table S6-S9 for full media descriptions and details.

Syringe pump configurations

Fresh media were infused through the microfluidic device by a syringe pump after cell loading. Programmable syringe pumps (NewEra pump systems, SKU:4000) were used in mother machine microfluidic experiments. In steady-state experiments, the syringes were programmed to continuously pump the growth media at a rate of 0.8 mL/hour for each main trench throughout the experiment. In nutrient up-shift and down-shift experiments, the syringes were programmed to switch between different media for each main trench. See Fig. 3, Fig. S1, and Tables SII-SIX for full experimental settings and details.

Microscopy and image acquisition

We performed time-lapse phase-contrast and fluorescence imaging on a Nikon ECLIPSE Ti2 inverted microscope equipped with a 100x oil-immersion objective (PH3, numerical aperture = 1.45) and the ORCA-Fusion BT Digital CMOS camera (C15440-20UP). Phase-contrast imaging was performed with an exposure time of 200 ms, and msfGFP channels were imaged at 20 ms using excitation wavelengths of 488 nm (Spectra III, lumencor) at 5% transmission, and mCherry channels were imaged at 10 ms using excitation wavelengths of 594 nm at 10% transmission. Phase-contrast images were captured every 4 minutes. The physiological effect of transmitted light on cell growth is negligible. To avoid cell photobleaching and phototoxicity due to the fluorescence illumination, msfGFP fluorescence images were captured every 20 minutes, and mCherry images were captured every 2 hours.

#### 3. Image processing

Image processing with the MM3 image analysis pipeline

Imaging processing follows the previously developed Python-based mm3 software pipeline (Thiermann et al., 2024). The MM3 image analysis pipeline includes channel compilation and designation, background subtraction, cell segmentation, cell tracking through lineages, and data output and analysis. Single-cell physical properties, such as length, width, and growth rates, are calculated through lineages.

Measurement of single-cell physiological quantities

Instantaneous single-cell growth rate for *E. coli* is calculated by measuring how fast the cell length increases over time as follows: (1) After image processing, the cell length $L(t)$ at time $t$ is collected. (2) Exponential cell growth can be described as $L(t) = L_0 e^{\mu t}$, where $L_0$ is the initial cell length. Take the natural log of $L(t)$, we have $lnL(t) = lnL_0 + \mu t$, where μ is the cell growth rate, estimated from the slope using linear regression. We calculated the instantaneous single-cell growth rates by fitting μ to the increment of cell length data over five consecutive time frames in an image series, corresponding to a 16-minute time interval. We used this specific number of frames because fewer frames would increase noise in transient growth rate estimates, while more frames would reduce temporal resolution and obscure rapid changes. (3) Population-level doubling time is calculated from the growth rate by $doubling\ time = ln(2)/\mu$, where μ is the average of the instantaneous single-cell growth rate, a proxy of volume expansion rate as cell width is mostly constant all the time in a steady state condition.

Single-cell fluorescence per volume ($\mu m^3$), i.e., the fluorescence concentration, is also quantified from image series. As the fluorescence images were captured less frequently than the phase contrast images, there were zero values in the time series of fluorescence concentration. To minimize discontinuities and maintain local trends before calculating the instantaneous fluorescence rates, zeros at both ends of the time series are replaced with the nearest non-zero values, and zeros between non-zero elements are replaced with the preceding non-zero values.

<u>Quantification of the surplus</u>

Surplus is quantified as the fraction of the total amount of gene product that is not contributing to the cell's steady-state growth rate. To obtain this quantity, a three-step analysis was performed as follows: (1) Combine steady state experiments for a specified growth medium with various arabinose concentrations for corresponding parent and tCRISPRi cells, and calculate the instantaneous growth rates and fluorescence concentrations. These growth rate and fluorescence intensity data were then normalized to the median of the parent cells' instantaneous growth rates and fluorescence concentrations, respectively. The normalized growth-rate versus fluorescence-concentration data were plotted as scatter plots in Figs. 1D and 2C. (2) Then the growth rates versus fluorescence concentration scatter points were fitted with a Hill function as $\mu = \frac{p^n}{K^n + p^n}$, where μ is the normalized growth rate, with unit $hour^{-1}$. $p$ represents the normalized target gene expression level (equal to the normalized fluorescence concentration), measured from the transcriptional or translational reporter at the same time point from the same cell, with an arbitrary unit (a.u.). $K$ is the normalized target gene expression level where the normalized growth rate is 0.5, similar to the Michaelis-Menten constant. And $n$ is the Hill coefficient. (3) After fitting, the normalized gene expression level fraction corresponding to above 90% of the normalized growth rate is quantified as the surplus, as shown in Fig. 2. Using a different threshold for normalized growth rate changes the surplus quantity slightly but does not alter the overall trend in the surplus of essential genes across different growth media (Fig. 2B, Fig. S2B and Fig. S4).

An alternative method to quantify surplus without fitting is also discussed in this study. With scatter plot data obtained in step (1) above, we can also quantify surplus using the rolling median curve, where the normalized fluorescence intensity fraction above 90% of the normalized growth rate is quantified as the surplus, as shown in Fig. S2A. This method yields a similar trend and conclusion regarding surplus across various growth media (Fig. 2B, Fig. S4).

Tracking of the recoverable cells post media up-shift

In up-shift experiments, we observed that some cells failed to adapt to rapid growth after being shifted to a fast-growth medium, shown as gray dots in Figs. 3B and S7B. To focus on cells that can eventually recover, we applied a lineage classification approach. First, complete cell lineages were reconstructed as a directed graph, linking daughter cells to their respective mother cells. Next, we identified the maximum independent subsets of these lineages within the graph. Finally, a threshold-based method was used to classify lineages according to their temporal growth dynamics: those achieving growth recovery following the media shift were labeled as “fast,” while the remainder were classified as “slow”. Figs. 3B and S7B depict fast lineages as green or red dots and slow lineages as gray dots.

Calculation of the adaptation time

To obtain the adaptation rate $k$ during a nutrient up-shift, we fitted the post-shift growth rates with an exponentially asymptotical curve $\mu(t) = - Ae^{-kt} + B$, where $\mu$ is the growth rate during adaptation, with unit $hour^{-1}$, and $B$ is the maximum steady-state growth rate after adaptation. $B - A$ defines the minimum steady-state growth rate at the beginning of the media shift, where $t = 0$. Both $A$ and $B$ can be extracted from the median growth rates in pre- or post-media shift data. The adaptation rate $k$ is the only fitted parameter, and its dependence on surplus for different gene is shown in Fig. S7C.

The adaptation time $\tau$ is defined as the time duration after the shift when the growth rate first reaches 90% of the maximum growth rate along the fitted asymptotical curve $\mu(t)$. The average adaptation time across all single-cell data is shown in green or red shades in Fig. 3B i. and iii., and in Figure S7B i. and iii. The single-cell adaptation time was obtained via similar steps, yet by fitting the $\mu(t)$ curve for individual cell lineages (Fig. 3B ii. and Fig. S7B ii.).

Analysis of the cell death probability

During the essential gene repression experiment, the main phenotypes of cell death in E. coli we observed include lysis, filamentation, and growth arrest. To quantify the likelihood of each cell death phenotype under different perturbation conditions of the surplus, we developed a multiparametric analysis pipeline that combined manual annotation with morphology-based feature extraction.

Cell filamentation leads to eventual cell death. We used threshold-based image analysis to determine if a cell becomes filamentous. Specifically, a cell is classified as filamentous when its length exceeds twice the average length of parent cells before division.

Cell lysis was manually annotated for mother cells. Lysis was identified based on visible morphological collapse, including disappearance of the defined cell boundary, and rapid dispersion or leakage of intracellular material relative to their previous frame. We also developed a graphical user interface (GUI) using Python and napari viewer for recording cell lysis events in the analysis.

Dependence of cell death probability on gene expression was quantified as follows. For each mother-cell lineage, cell fate was classified as normal growth, lysis, growth arrest or filamentation. For lineages that eventually exhibited a death-associated phenotype (filamentation or lysis), fluorescence concentration was quantified prior to the event: in steady-state conditions, values were averaged over the 1 h interval immediately preceding the onset of detectable death phenotype; in nutrient upshift experiments, fluorescence was defined from the interval immediately before the upshift; and in starvation–recovery experiments, it was measured immediately prior to starvation, enabling consistent comparison of initial conditions. Lineages were then ranked by fluorescence intensity and partitioned into bins containing equal numbers of cells (typically 45, or 60 where indicated). For each bin, the death probability was computed as the fraction of lineages that ultimately underwent filamentation or lysis divided by the total number of lineages in that bin. This nonparametric binning procedure yields an empirical estimate of death probability as a function of gene expression level while maintaining sufficient sampling within each bin for statistical robustness.
Growth-arrest analysis was also performed. Growth arrest is defined as cells that maintain a near-zero growth rate for approximately three or more cell-cycle durations before the end of imaging. [mention the growth arrest probability is calculated in a similar way to the cell death analysis]

**Title:** Emergent, cost-free surplus of core biosynthesis governs bacterial fitness

#### Supplemental Figures

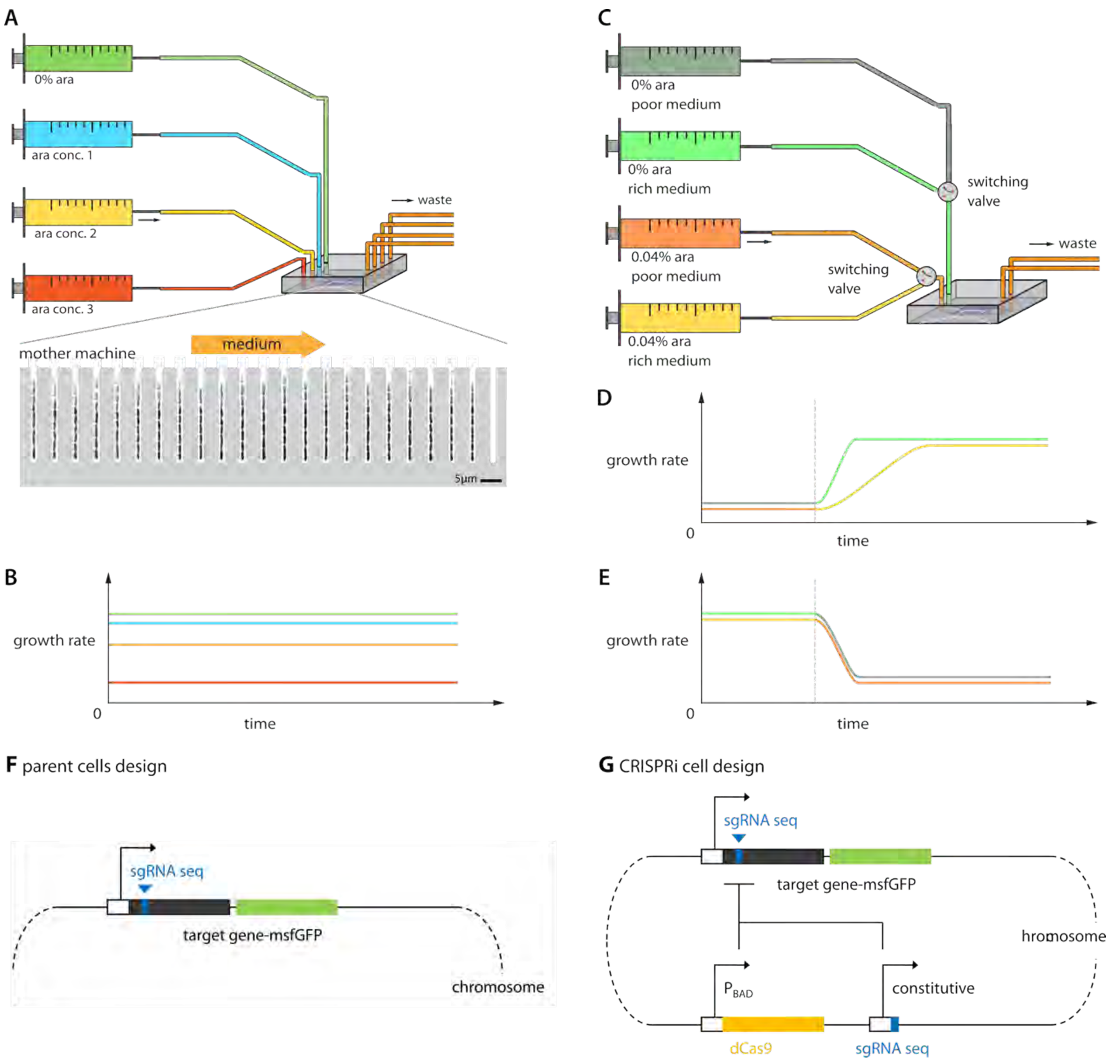

**Figure S1: Illustrations of experimental setup. A.** Configuration of experimental conditions for assessing steady-state cell growth under varying levels of surplus. A multiplex mother machine with four independent devices is used here. **B.** Visualization of growth rates during steady-state conditions. Colored lines correspond to different growth conditions. **C.** Setup for nutrient shift experiments. The syringe pump is programmed to switch between poor and rich media, with or without the inducer arabinose (are) to repress the gene of interest. **D.** Illustration of predicted growth rate adaptation following a nutrient up-shift. The two lines represent the adaptation with and without gene repression; the same for **E.**, which is the illustration of predicted growth-rate adaptation following a nutrient down-shift. **F.** Parent strains are engineered to express the superfolder GFP following the target gene as a transcriptional or translational reporter, enabling the quantification of gene expression level. **G.** The design of the CRISPRi strains. The genetic construction details are summarized in Methods.

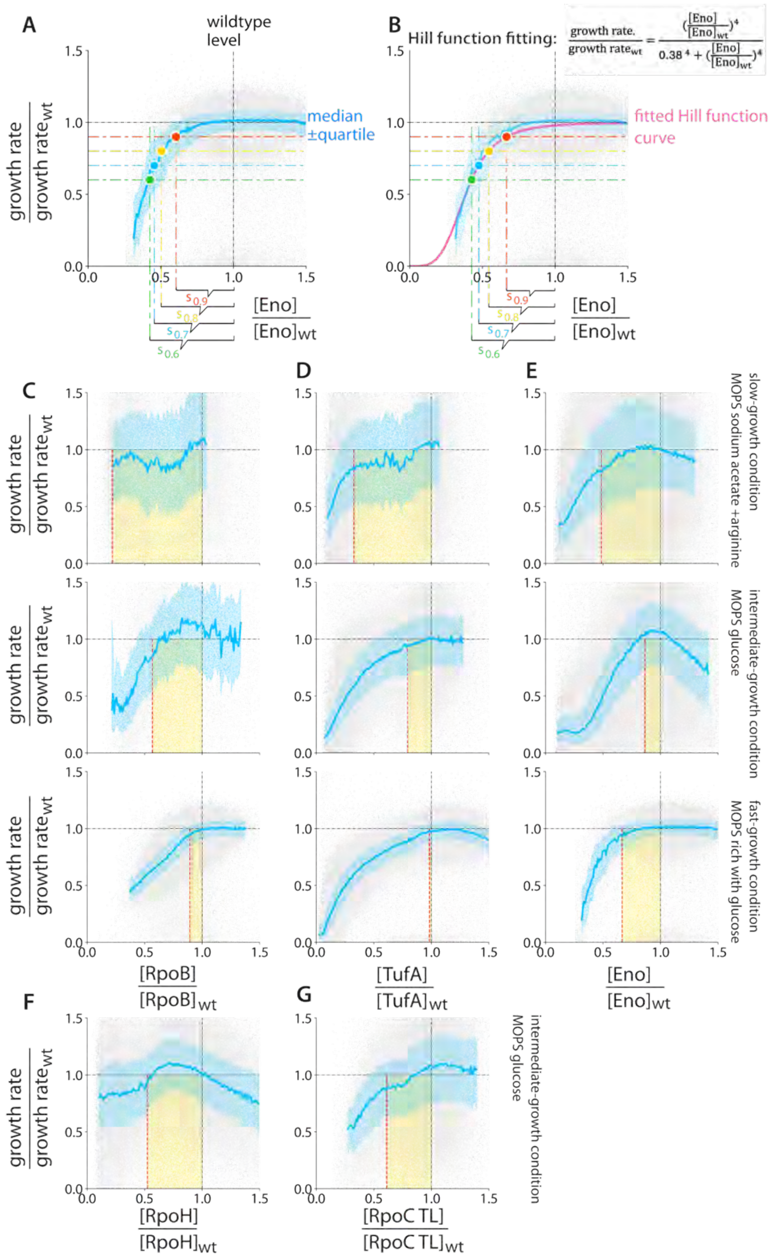

**Figure S2. Quantifying surplus for central biosynthetic components. A.** Surplus quantification via plateau detection using the rolling median of growth vs. expression data. Expression level of enolase, [Eno], is shown here. Each horizontal dashed line represents a threshold as a fraction of the wild-type growth rate. An expression level exceeding the threshold is considered a surplus. Surpluses $s_{0.6}$, $s_{0.7}$, $s_{0.8}$, and $s_{0.9}$, quantified with different thresholds of 0.6, 0.7, 0.8, and 0.9, respectively, are shown. **B.** Similar surplus quantification via plateau detection in the fitted Hill-function curve of growth vs. expression data. The Hill coefficient was fitted to be 4. **C-G.** The scatter plot (light gray) shows single-cell growth rate versus *rpoB*, *tufA,* and *eno* expression levels across different media, corresponding to the data points shown in Figure 2C. $s_{0.9}$ determined by Hill-function fitting is used here, similar to Figs. 1 and 2. The blue line represents the median growth rate, while the shaded blue area indicates the interquartile range. Surplus for *rpoB*, *tufA,* and *eno* in each condition is shaded in yellow.

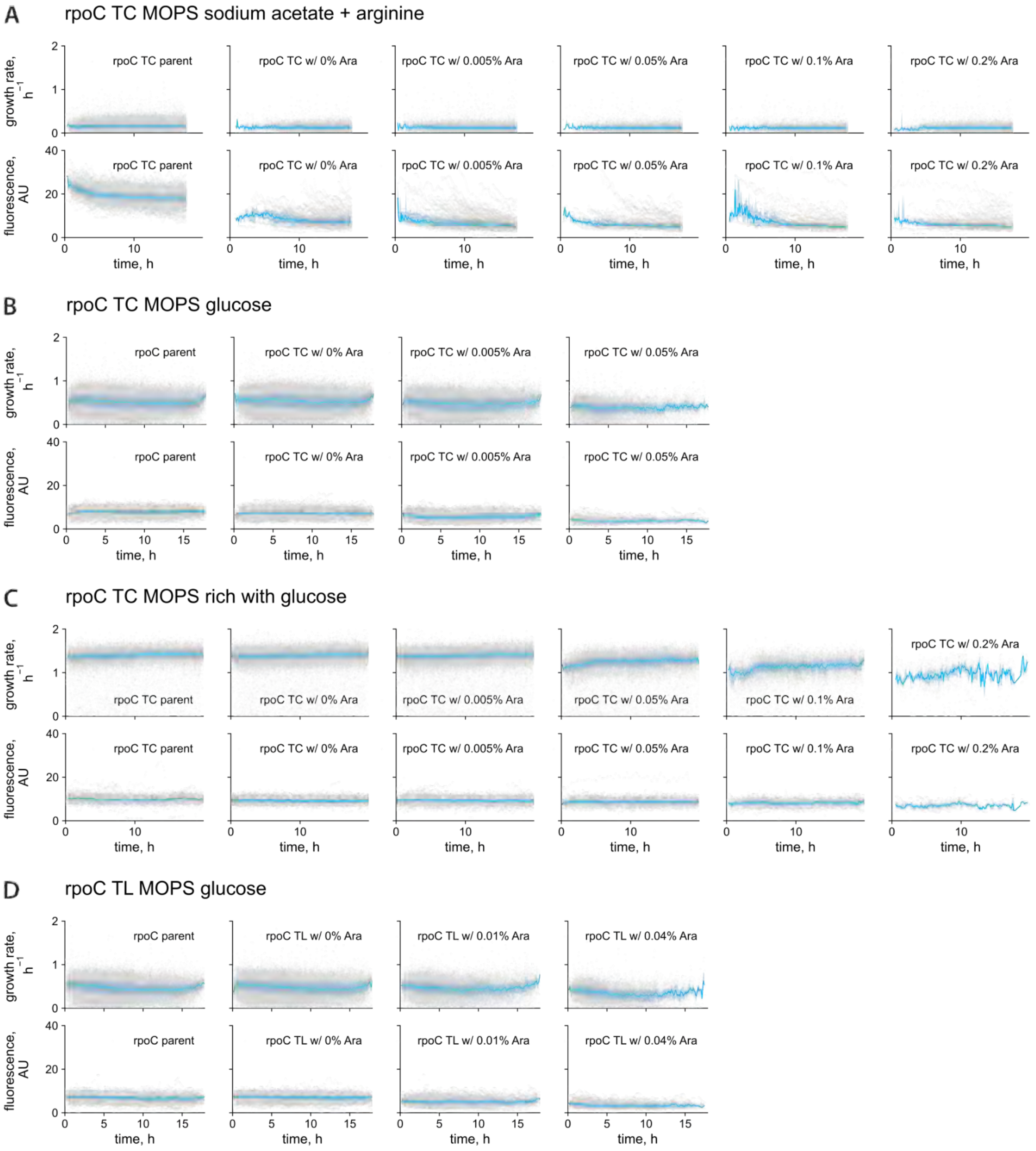

**Figure S3. Growth-rate time course and fluorescent-intensity time course for steady-state growth of *rpoC* cells with transcriptional (TC) or translational (TL) reporter in different growth media.** The scatter plot (light gray) shows single-cell growth rate versus time. The blue line represents the median growth rate, while the shaded blue area indicates the interquartile range.

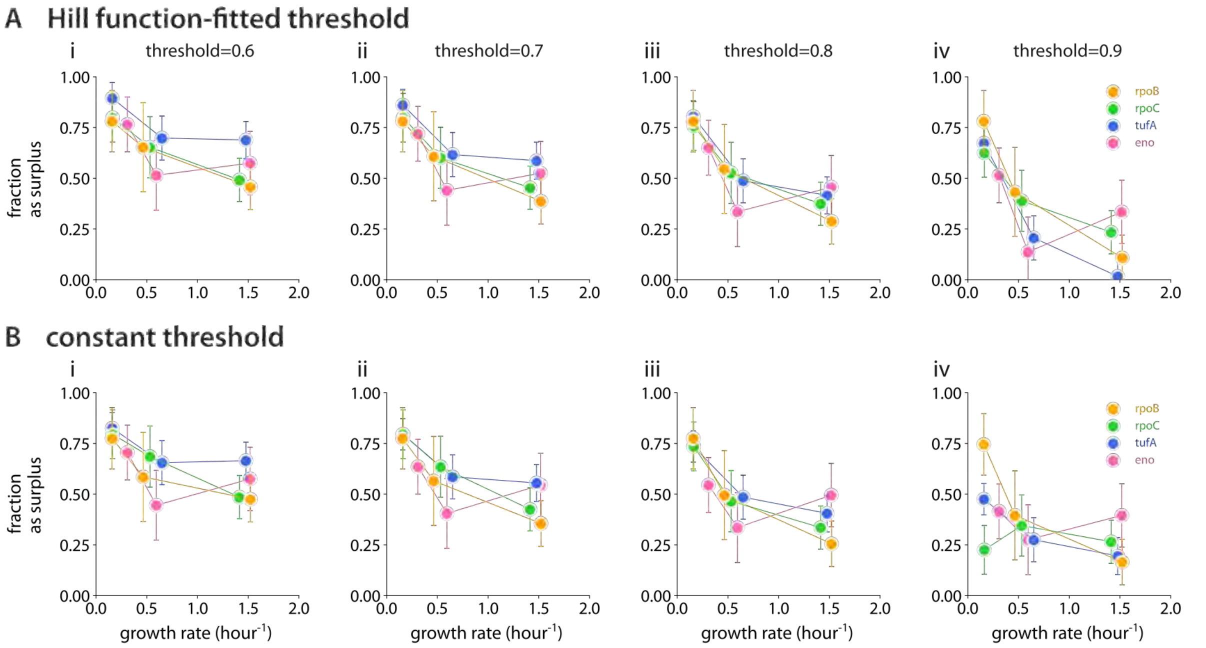

**Figure S4. Dependence of surplus on the growth rate in different nutrient media. A.** Surplus quantification via plateau detection in the fitted Hill function of growth vs. expression data. Plots i-iv show surplus quantified with different thresholds of 0.6, 0.7, 0.8, and 0.9, respectively. **B.** Surplus quantification via plateau detection in the rolling median of growth vs. expression data. Plots i-iv show surplus quantified with different thresholds of 0.6, 0.7, 0.8, and 0.9, respectively.

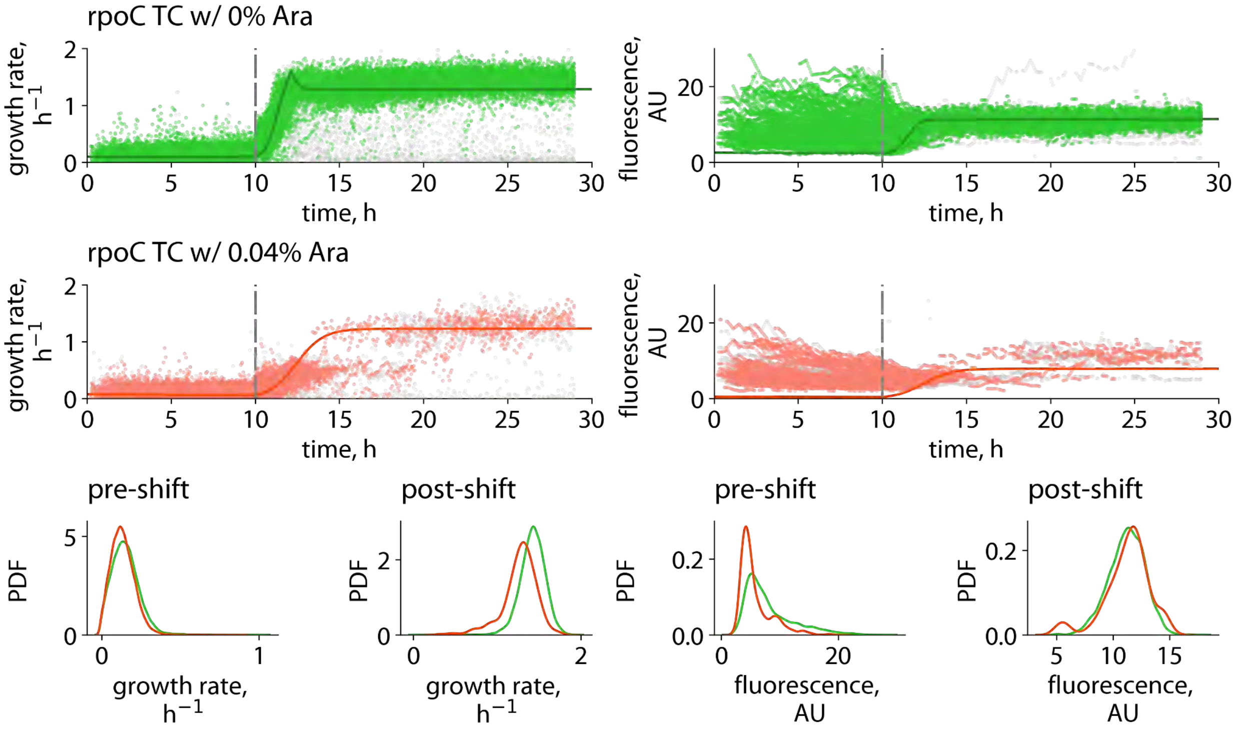

**Figure S5. Nutrient up-shift experiments with rpoC cells show delays in adaptation when RpoC surplus is repressed.** In the up-shift experiments, cells were initially grown for 10 hours at steady state in a slow-growth medium (MOPS supplemented with sodium acetate and arginine), followed by an immediate switch to a fast-growth medium (glucose-rich MOPS medium). **A.** Time-course profiles of (i) growth rate ($h^{-1}$) and (ii) fluorescence intensity (AU) for rpoC (transcriptional reporter, TC) cells cultured without arabinose (Ara, 0%) are shown. Green scatter points represent single-cell data from adapting cells, whereas gray points indicate cells that did not adapt to the enriched medium. Dark green lines represent model fits. Details of the model and parameter values are provided in the Supplemental Information and Table S1. **B.** Time-course profiles of (i) growth rate ($h^{-1}$) and (ii) fluorescence intensity (AU) for cells cultured with 0.04% arabinose. Red scatter points indicate adapting cells, whereas gray points represent non-adapting cells. Dark red lines show model fits. Plot styles are the same as in panel A. **C.** Probability density functions (PDFs) of single-cell growth rates before and after the nutrient shift, estimated using Gaussian kernel density estimation. Green curves correspond to 0% Ara and red curves to 0.04% Ara. **D.** PDFs of single-cell fluorescence intensities before and after the nutrient shift. Plot styles are the same as in panel C.

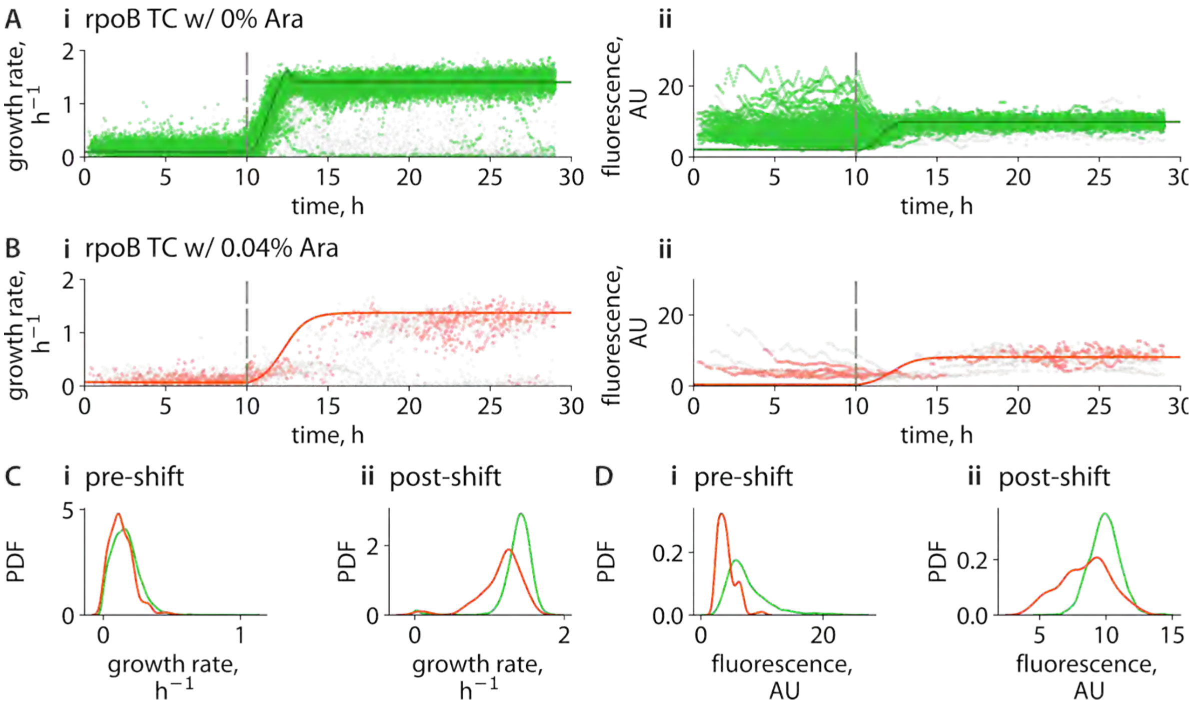

**Figure S6. Nutrient up-shift experiments with *rpoB* cells show delays in adaptation when RpoB surplus is repressed.** In the up-shift experiments, cells were initially grown for 10 hours at steady state in a slow-growth medium (MOPS supplemented with sodium acetate and arginine), followed by an immediate switch to a fast-growth medium (glucose-rich MOPS medium). **A.** Time-course profiles of (i) growth rate ($h^{-1}$) and (ii) fluorescence intensity (AU) for rpoC (transcriptional reporter, TC) cells cultured without arabinose (Ara, 0%) are shown. Green scatter points represent single-cell data from adapting cells, whereas gray points indicate cells that did not adapt to the enriched medium. Dark green lines represent model fits. Details of the model and parameter values are provided in the Supplemental Information and Table S1. **B.** Time-course profiles of (i) growth rate ($h^{-1}$) and (ii) fluorescence intensity (AU) for cells cultured with 0.04% arabinose. Red scatter points indicate adapting cells, whereas gray points represent non-adapting cells. Dark red lines show model fits. Plot styles are the same as in panel A. **C.** Probability density functions (PDFs) of single-cell growth rates before and after the nutrient shift, estimated using Gaussian kernel density estimation. Green curves correspond to 0% Ara and red curves to 0.04% Ara. **D.** PDFs of single-cell fluorescence intensities before and after the nutrient shift. Plot styles are the same as in panel C.

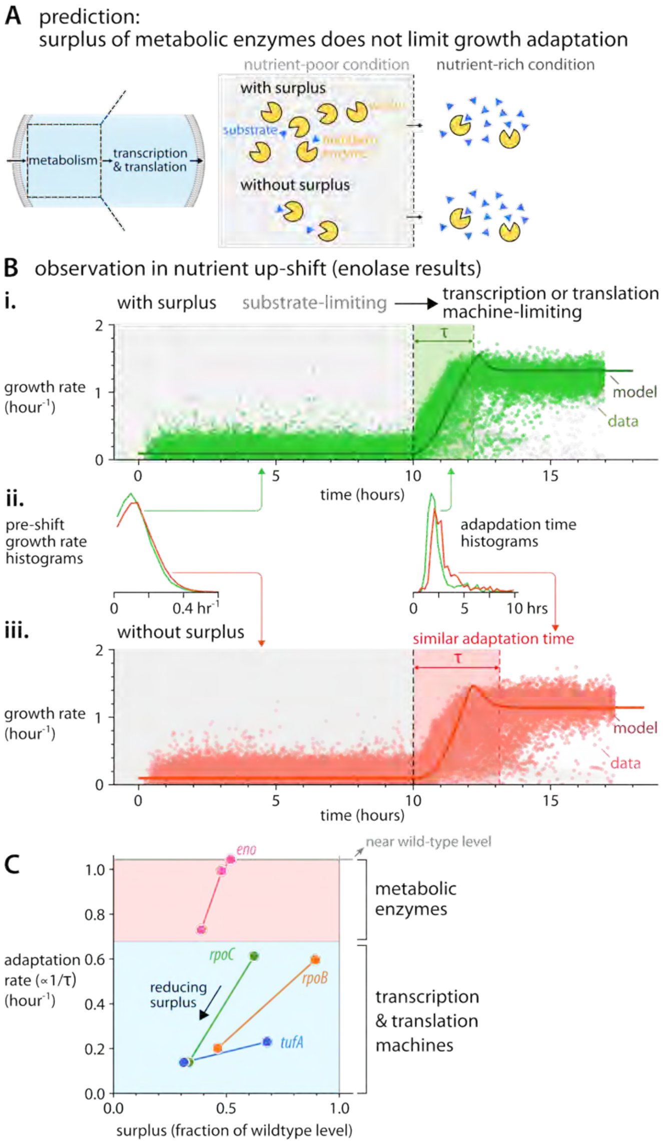

**Figure S7. The surplus of metabolic enzymes has little effect on growth adaptation rate during a nutrient up-shift. A.** An illustration showing that the surplus of metabolic enzymes may not limit biosynthesis during a nutrient up-shift. In nutrient-poor conditions, substrate scarcity limits growth, whereas in nutrient-rich conditions, downstream transcription and translation machinery limit growth. **B.** Time-course plots showing that the growth adaptation time is similar with (**i.**) or without (**iii.**) a surplus of enolase when shifting from MOPS medium with glucose and arginine to MOPS-rich medium with glucose [see other growth conditions in Fig. S17]. Different from RpoC, enolase's histograms of pre-shift growth rate and the adaptation time remain unchanged with or without surplus (**ii.** histograms in the middle). Solid lines represent the model calculation with the enolase surplus being the only free parameter [SI]. Data representation is similar to Fig. 3B. Light gray data points represent non-growing cells excluded from the adaptation analysis, and more non-growing cells are observed when enolase surplus is repressed. **C.** The adaptation rate, proportional to the inverse of the adaptation time, decreases when the surplus of transcription or translation machines is reduced, but is less dependent on the surplus of metabolic enzymes such as enolase.

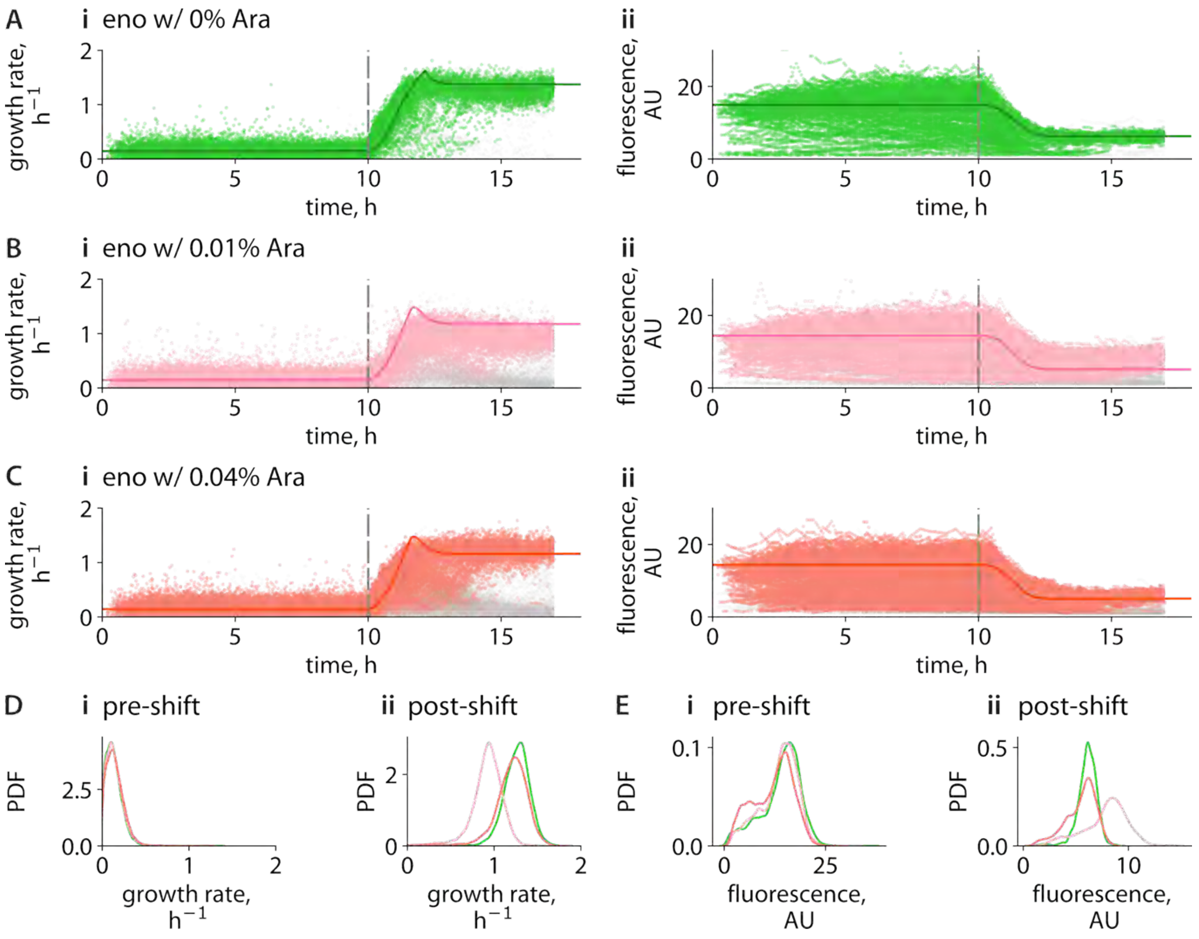

**Figure S8. Nutrient up-shift experiments with *eno* cells when enolase surplus is repressed.** In the upshift experiments, cells were initially grown for 10 hours at steady state in a slow-growth medium (MOPS supplemented with glucose and arginine), followed by an immediate switch to a fast-growth medium (MOPS rich with glucose). **A-C.** Time-course profiles of (i) growth rate (hour$^{-1}$) and (ii) fluorescence intensity in arbitrary units (AU) for *eno* (transcriptional reporter) cells. **A.** Up-shift experiment time course with no arabinose (Ara, 0%). **B.** Up-shift experiment time course with 0.01% Ara. **C.** Up-shift experiment time course with 0.04% Ara. Colored scatters represent adapting cells; gray points indicate non-adapting cells. **D**. PDFs of single-cell growth rates before (i) and after (ii) the nutrient shift. **E.** PDFs of single-cell fluorescence intensities before (i) and after (ii) the shift. Color codes are consistent with **A-C**.

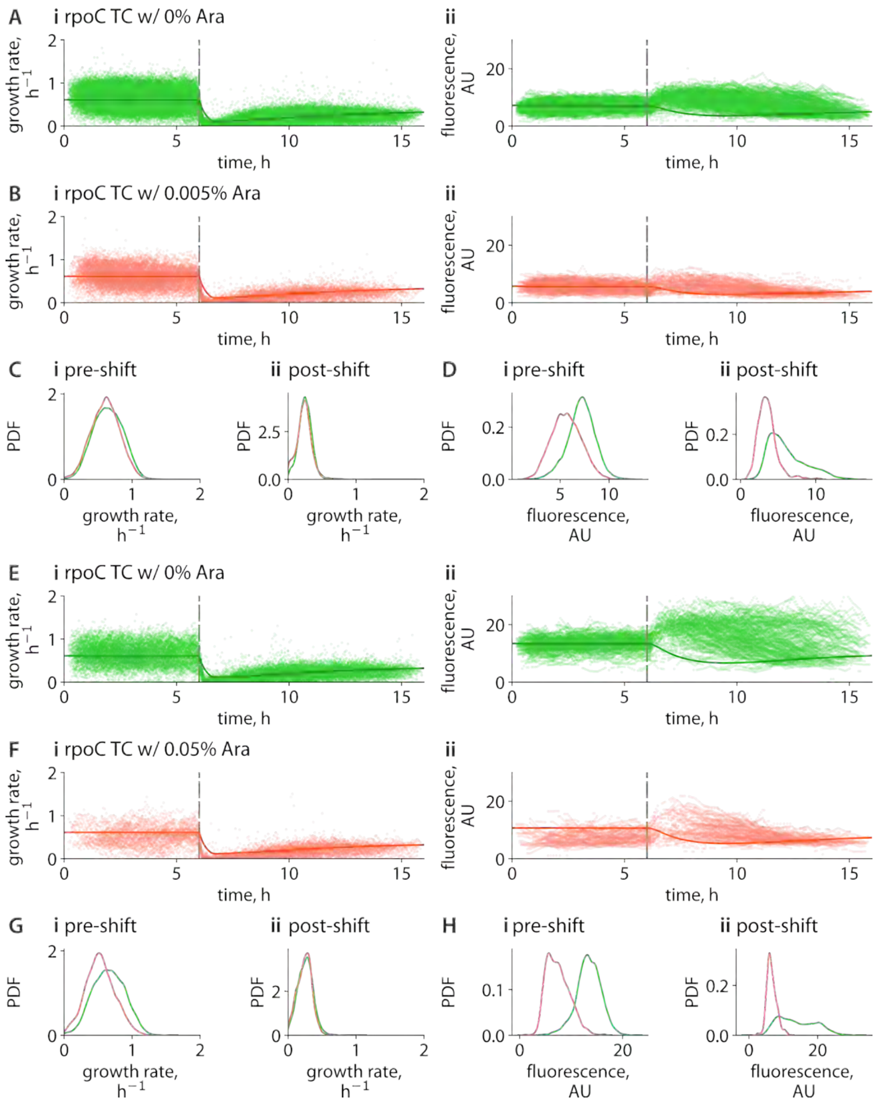

**Figure S9. Nutrient down-shift experiments with *rpoC* cells when RpoC surplus is repressed.** In the down-shift experiments, cells were initially grown for 6 hours at steady state in a fast-growth medium (MOPS supplemented with 11 amino acids and glucose), followed by an immediate switch to a slow-growth medium (MOPS with sodium acetate). A-D and E-H are two sets of experiments. **A-B.** Time-course profiles of (i) growth rate (hour$^{-1}$) and (ii) fluorescence intensity in arbitrary units (AU) for *rpoC* cells. **A.** Down-shift experiment time course with no arabinose. **B.** Down-shift experiment time course with 0.005% Ara. Plot styles are the same as **A**. **C**. PDFs of single-cell growth rates before (i) and after (ii) the nutrient shift, with green line for 0% Ara and red for 0.005% Ara. **D.** PDFs of single-cell fluorescence intensities before (i) and after (ii) the shift. Plot styles are the same as **C**. **E-F.** Time-course profiles of (i) growth rate (hour$^{-1}$) and (ii) fluorescence intensity in arbitrary units (AU) for *rpoC* cells. **E.** Down-shift experiment time course with no arabinose. **F.** Down-shift experiment time course with 0.05% Ara. Plot styles are the same as **E**. **G**. PDFs of single-cell growth rates before (i) and after (ii) the nutrient shift, with green line for 0% Ara and red for 0.05% Ara. **H.** PDFs of single-cell fluorescence intensities before (i) and after (ii) the shift. Plot styles are the same as **G**.

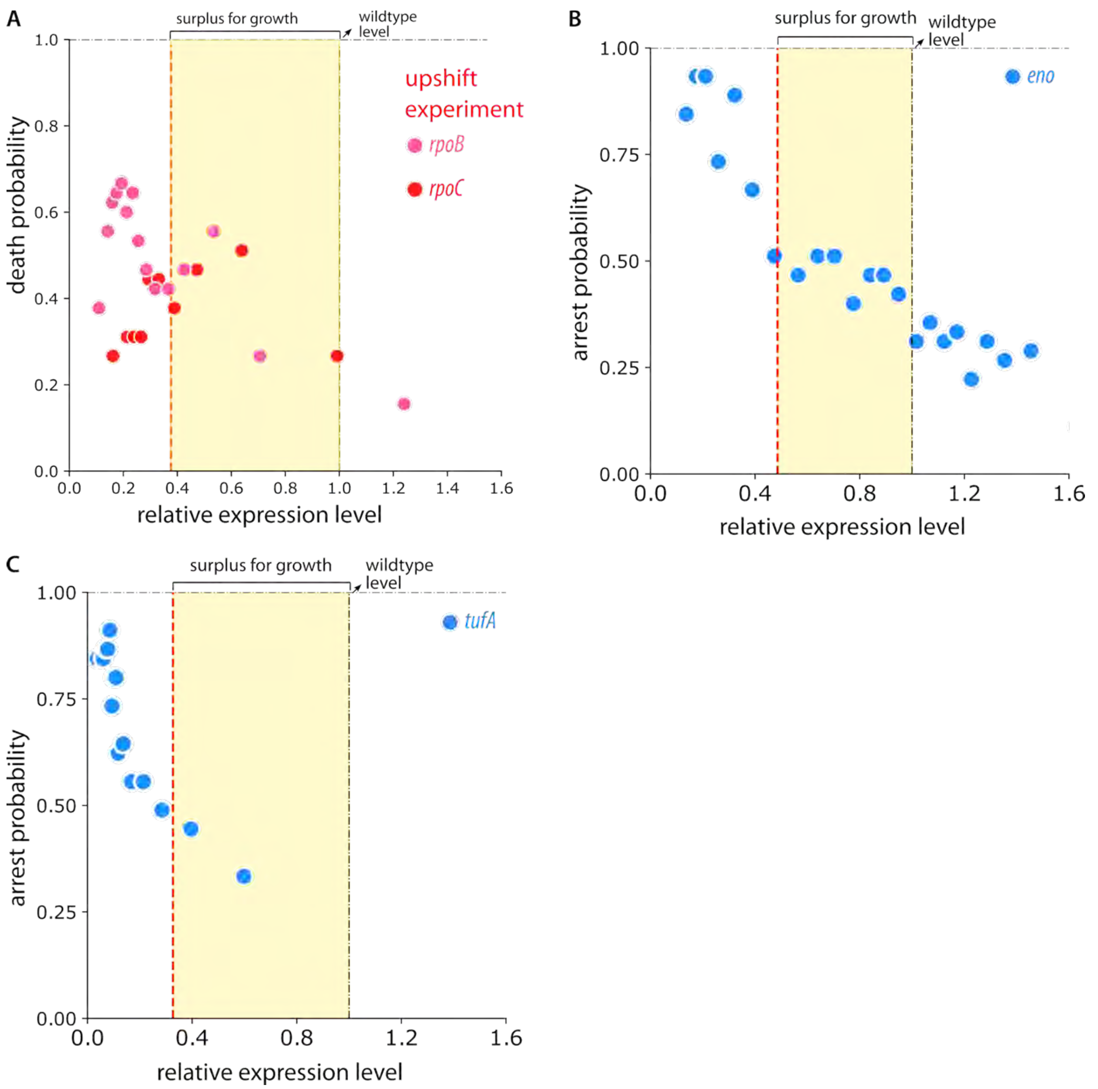

**Figure S10. Cell fates in nutrient transitions. A.** In upshift experiments, transcriptional responses of suppressing surplus of RNA polymerase components, RpoB and RpoC, exhibit a more pronounced increase in cell death probability than observed under steady-state conditions. **B.** Scatter plot illustrates the relationship between relative cell arrest probability and relative *eno* expression level in the nutrient upshift experiment. In the upshift experiments, cells were initially grown for 10 hours at steady state in a slow-growth medium (MOPS supplemented with sodium acetate and arginine), followed by an immediate switch to a fast-growth medium (MOPS rich with glucose). Surplus for *eno* in the preshift condition is shaded in yellow. Cells without surplus of enolase show excessive growth arrest following nutrient upshift. **C.** Same analysis as in **B**, for the nutrient upshift for tufA cell.

### Supplemental information for "Emergent, cost-free surplus of core biosynthesis governs bacterial fitness."

Huijing Wang[1], Dotan Goberman[2], Christopher Aldrich[3], Rami Pugatch[2,*], Fangwei Si[1,3,*]

*1. Department of Physics, Carnegie Mellon University*

*2. Department of Industrial Engineering and Management, Ben-Gurion University of the Negev*

*3. Department of Biomedical Engineering, Carnegie Mellon University*

**CONTENTS**

I. Supplemental notes: Modeling the surplus of central biosynthetic components and its effect on growth adaptation 2

A. Model description 2

B. Basic assumptions of the model 4

C. Numerical calculation of the model 7

D. Comparison with other bacterial growth models 9

E. Depletion probability of core biosynthetic components within an autocatalytic network 14

F. Justification of course-grained modeling of growth dynamics 15

II. Supplemental tables 16

III. Supplemental figures 31

References 42

#### I. SUPPLEMENTAL NOTES: MODELING THE SURPLUS OF CENTRAL BIOSYNTHETIC COMPONENTS AND ITS EFFECT ON GROWTH ADAPTATION

##### A. Model description

We use a simplified model, presented in [1], which agglomerates the transcription-translation machinery, including RNA polymerase (RNAP) and ribosome with accessories, into a single unit, $U$, capable of self-replication and synthesizing any other protein in the cell (Eqs. S1, S2). In particular, the protein $P$ in the model represents agglomerated metabolic proteins that feed $U$ with building blocks and energy for protein synthesis, annotated by $F$, which drives the process (Eq. S3). The $P$s themselves are fed by the raw materials (nutrient molecules), $f$. $f$ is accumulated in proportion to total biomass, $U + P$, representing the case that each cell is always surrounded by the same concentration of nutrient molecules in a constant environment. The number fraction of $U$s allocated to synthesize more $U$s, $\alpha$, is a control parameter that is a function of the state (Eq. S5). Note that $\alpha$ is bound from zero to $\phi_{\max}$, where $\phi_{\max}$ is the maximal number fraction of ribosomes [2]. Here, we use $U$, $P$, $F$, and $f$ to represent the absolute number of each biomolecule for simplicity, instead of the absolute mass often used in other models, which we will show does not affect the model behaviors or conclusions.

To control $\alpha$, we employ perhaps the simplest model, which aims to lock $\alpha$ so that the ratio between substrate supply $F$ and the number of $U$s is constrained within a certain range around a set point of $\frac{U}{F}$, $S_0$, which is a positive number not much smaller than 1. This ensures that there is neither too much excess nor a shortage of resources to feed $U$, and locking $\alpha$ around a reasonable range between 0 and $\phi_{\max}$. A negative term, $-k_1\alpha$, is added to ensure that $\alpha$ converges around the default values, forming a negative feedback loop. If the set point of $\frac{U}{F}$, $S_0$, is less than 1, there is a shortage of resources to feed $U$, which means there is a surplus of $U$. If $S_0$ is greater than 1, there is an excess of resources to feed $U$, which means that there is a surplus of $F$. The equation of $\frac{d\alpha}{dt}$ resembles a partial PID controller with the first term on the right-hand side being an integral gain that corrects $\alpha$ based on the difference between $S_0$ and $\frac{U}{F}$, and the second term being an internal decay to constrain the variation in $\alpha$. Other forms of control equations that impose negative feedback on $\alpha$ based on the states of $F$ and $U$ could yield similar model behavior and conclusions. See more discussions in Suppl. Notes I D and more rationale in Suppl. Notes I F).

The full set of equations for the model is as follows.

$$\frac{dU}{dt} = \frac{\alpha \cdot \min(U, F)}{\tau_U}, \tag{S1}$$

$$\frac{dP}{dt} = \frac{(\phi_{\max} - \alpha) \cdot \min(U, F)}{\tau_P}, \tag{S2}$$

$$\frac{dF}{dt} = \frac{\min(P, f)}{\tau_F} - \frac{\alpha \cdot \min(U, F)}{\tau_U} - \frac{(\phi_{\max} - \alpha) \cdot \min(U, F)}{\tau_P}, \tag{S3}$$

$$\frac{df}{dt} = k \cdot \frac{d(U + P)}{dt} - \frac{\min(P, f)}{\tau_F}, \tag{S4}$$

$$\frac{d\alpha}{dt} = k_2 \left(S_0 - \frac{U}{F}\right) - k_1\alpha, \quad \text{with the constraint, } 0 \leqslant \alpha \leqslant \phi_{\max}, \tag{S5}$$

where $\tau_U$, $\tau_P$ and $\tau_F$ are the time scales to produce one $U$, $P$, and $F$, respectively, $k_1$ and $k_2$ are the control rates for $\alpha$, and $k$ represents the proportionality between raw materials (nutrient molecules) and total biomass $U + P$ (related to the quality of nutrients and the characterization of the nutrient uptake process). The representative numerical values and physical units for each parameter used in our calculation are shown in Table SI. The illustration of the model is shown in Fig. N1.

##### B. Basic assumptions of the model

The introduction of minimum functions distinguishes our model from others. Although explained in the original work [1], we reiterate its meaning here. We use Eq. S1 to describe protein translation as an example. $U$ represents the number of ribosomes, with $F$ being the number of amino acids, $\tau_U$ the time to produce all ribosomal proteins and assemble them into a new ribosome, and $\alpha$ the number fraction of ribosomes that produce ribosomal proteins. When amino acids (ribosome substrates) are abundant enough in the cell, e.g., when cells grow in a nutrient-rich condition, at a given moment, they could outnumber ribosomes, i.e., $U \leqslant F$. In this case, all ribosomes are saturated with amino acids, and the protein production rate is subject to the number of ribosomes, not the number of amino acids, a portion of which is free and not used for translation. In contrast, when the supply of amino acids is limited in the cell, e.g., when cells grow under nutrient-poor conditions, at a given moment, the substrate molecules could be fewer ribosomes, i.e., $F < U$. In this case, only a portion of the ribosomes bind to amino acids, and the protein production rate is subject to the number of ribosomes that bind to amino acids, which is equal to the number of amino acids that bind to ribosomes. The same reasoning applies to the production of non-ribosomal proteins, $P$, from $F$ and the substrate, $F$, from $f$.

In our model, the time scales for protein production, $U$ or $P$, and for building blocks, $F$, are invariant; that is, $\tau_U$, $\tau_P$, and $\tau_F$ are constants. This assumption seems inconsistent with experimental evidence that some of these time scales are condition-dependent, e.g., the slowdown of translation elongation rate in nutrient-poor conditions [3]. However, in our model, using the production of $U$ as an example, excessive $U$ would increase under nutrient-poor conditions due to the scarcity of $f$ and $F$, i.e. $U > F$. Therefore, in such a case, when calculating the average time scale over the total population of $U$, the equivalent time scale is $\frac{F}{U}\tau_U$. Across nutrient conditions, the equivalent time scale of $U$ production is $\frac{\min(\mathrm{U,F})}{U}\tau_U$, which shows a bilinear trend against the nutrient-imposed steady-state growth rate, mimicking the observed nutrient-dependent translation elongation rate.

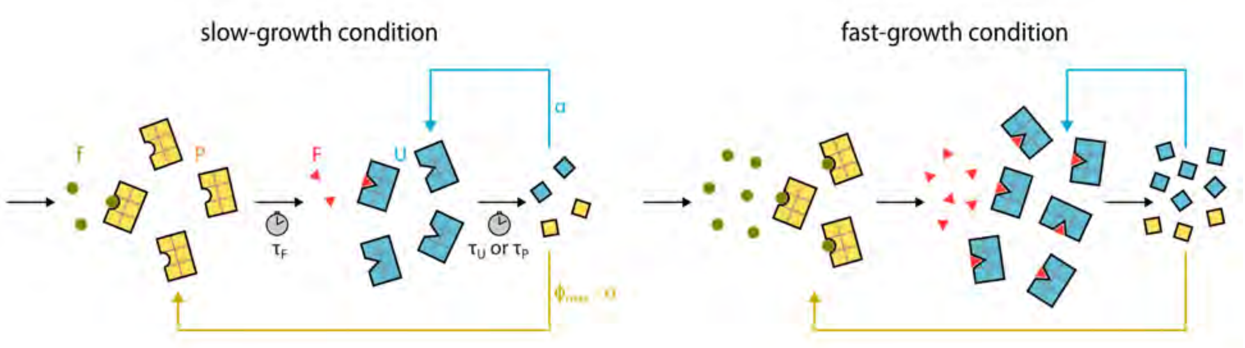

**Fig. N**1: **Model illustration.** **(A)** In slow-growth conditions, the raw materials $f$ are more scarce than its catalyst $P$, so are the building blocks $F$ versus its catalyst $U$. Therefore, $P$ and/or $F$ are the limiting factors for the respective biosynthetic processes. **(B)** By contrast, in fast-growth conditions, both $P$ and $U$ are outnumbered and saturated by their substrates. In this case. $P$ and/or $U$ are the limiting factors for the respective biosynthetic processes.

Note that the definition of enzyme surplus, e.g., for $U$, surplus $= \frac{U-\min(U,F)}{U}$, is not explicitly expressed in Eqs. S1-S5. Since in a steady state, $\frac{U}{F} \approx S_0$, we can show that the surplus is connected to the set point of $\frac{U}{F}$, $S_0$, by

$$\text{surplus} \approx \max(1 - \frac{1}{S_0}, 0). \tag{S6}$$

This means that, when $F \geqslant U$ in steady state, $S_0 \lessapprox 1$, and the surplus of $U$ is 0, and when $F < U$ in steady state, $S_0 \gtrapprox 1$, and the surplus is roughly equal to $1 - \frac{1}{S_0}$. Therefore, we can use $S_0$ as a single handle to control the production levels of $U$ and $P$, therefore tuning their surplus. Also note that this model is not for describing single-cell behaviors such as cell division and cell-size control, or the absolute mass of the major biomolecules, but just characterizing the absolute number of major biomolecules for simplicity, without losing the physical essence. Since the total protein mass is the majority of the total cell biomass, we use $U + P$ to approximate the total cell biomass, and therefore the growth rate $\mu$ can be defined as $\mu = \frac{d(U+P)}{dt}\frac{1}{U+P}$. The number fraction of $U$ and $P$ can be calculated as $\phi_U = \frac{U}{U+P}$, and $\phi_P = \frac{P}{U+P}$, which are closely related to their mass fractions. It can be shown that, when the system reaches a steady state, $\alpha = \phi_U$, and $1 - \alpha = \phi_P$. Using this highly simplified model allows us to capture and predict the steady-state growth rate and non-steady-state adaptive behavior of cells during nutrient shifts, with minimal assumptions, as we describe below in Suppl. Notes I C.

| **Fixed Parameter** | **Definition** | **Value** | **Unit** | **Notes** |
|---|---|---|---|---|
| $\tau_U$ | time of producing one $U$ | 9.6 | minute | [3, 4] |
| $\tau_P$ | time of producing one $P$ | 9.6 | minute | [4] |
| $\tau_F$ | time of producing one $F$ | 9.6 | minute | [1] |
| $\phi_{\max}$ | maximal number fraction of $U$ | 0.41 | unit-less | [2] |
| $k_1$ | control rate 1 for $\alpha$ | 1.2 | minute$^{-1}$ | [5]; fixed for up-shift data of this work |
| $k_2$ | control rate 2 for $\alpha$ | 0.1 | minute$^{-1}$ | [5]; fixed for up-shift data of this work |
| **Free Parameter** | **Definition** | **Value** | **Unit** | **Notes** |
| $k$ | proportionality between nutrient and biomass (related to nutrient quality) | between 1 and 2.5, depending on the growth media | unit-less | fixed from steady-state data of this work |

| | | | |
|---|---|---|---|
| $S_0$ | set point for $\frac{U}{F}$ | decreasing from around 4-5 to 1-3.5 when repressing *RpoC*, *RpoB*, or *tufA*; increasing from around 4-5 to 5-6 when repressing *eno* | unit-less |

**Table SI: Parameters used in the numerical calculation.**

##### C. Numerical calculation of the model

We numerically calculated the time-evolved solutions of Eqs. S1-S5 using an ODE solver (odeint) in Python. The majority of the parameters used and the numeric range of variables for the calculation can be fixed from the literature or from steady-state data in this work, as shown in Table SI [also see Figs. N2 and N3]. Among the fixed parameters, the time scale for transcription and translation, $\tau_U$ or $\tau_P$, is on the scale of minutes based on previous measurements [3, 4]. The time scale of the agglomerated chain of central metabolic reactions, $\tau_F$, is also on the scale of minutes [1]. The maximum number fraction of $U$, $\phi_{\max}$, can be estimated from the maximum ribosomal mass fraction [2]. The rate constants, $k_1$ and $k_2$, in the control equation for $\alpha$ are related to the time scale of regulation of ribosome synthesis via ppGpp ($\sim$ 1 minute$^{-1}$; see Suppl. Notes I D) [5]. Here, as phenomenological quantities, their values can be further constrained from the up-shift data of *rpoC* in this work [Fig. N4], and fixed for other up-shift data and the nutrient down-shift data, except for some fine-tuning of $k_1$ for the *eno* data to match the growth overshoot. [Figs. N5 and N6]. Note that their particular values do not affect the model behavior with respect to the role of surplus in cell adaptation to nutrient shifts. The nutrient quality-related parameter, $k$, is rather arbitrary, and undergoes a step function to stimulate the nutrient up-shifts or down-shifts. We fitted the model-calculated results to the experimental data by tuning the free parameters until we achieved a sufficiently small numerical difference. The specific procedure of numerical calculation is as follows.

First, we fixed most of the parameters based on the literature or steady-state data in this work, except for the rate constants $k_1$ and $k_2$. We then fitted the nutrient-quality-related parameter $k$ to the steady-state growth rate data. Note that a larger $k$ increases the growth rate monotonically. In steady-state cases, $\alpha$, is equal to the mass fraction of $U$, which is related to its number fraction, $\frac{U}{U+P}$, and all components, $U$, $P$, and $F$, are growing at the same rate as the biomass growth rate $\mu = \frac{d(U+P)}{dt}\frac{1}{U+P} = \frac{\alpha}{\tau_U}$, confirming the self-consistency of our model [Figs. N2 and N3].

Second, we applied the model with fixed parameters for the nutrient up-shift data of *rpoC* (considered as part of $U$) with and without gene suppression, to fit the set point for $\frac{U}{F}$, $S_0$, which is directly linked to the definition of surplus of $U$ by Eq. S6 [Fig. N4, model lines in Fig. 3B, Figs. S5-S6, Figs. N12-N14]. A tune-down of $U$'s surplus is equivalent to a decrease of $S_0$. A step function of the increase in $S_0$ is allowed to reflect the quick change in repression strength upon nutrient upshift, possibly due to catabolite repression on the CRISPRi system. We also fitted $k_1$ and $k_2$ to the overshoot behavior (damped oscillation) of the growth rate right after the up-shift, and fixed them for all other following conditions. Note that $S_0$ is fixed for unperturbed cells in all other nutrient up-shift and down-shift tests, including those of *rpoC*, *rpoB*, and *tufA* [Figs. N5 and N6, Figs. 3C, Figs. S5-S6, Figs. N12-N14]. Therefore, when testing the majority of our nutrient shift data, our model does not have free parameters, which demonstrates its predictive power.

We then applied the model with all the fixed parameters mentioned above to predict the nutrient down-shift data of *rpoC* with and without gene suppression [Fig. N5, Fig. 3C, Fig. S9, Fig. N16]. When calculating for the down-shift data, we allowed the nutrient molecule number $f$ to be depleted enough that the internal building block $F$ is also depleted, causing the growth rate to crash. We also applied the model for the up-shift data of *eno* (which can be assumed as part of $P$), without any free parameters (some fine-tuning of $k_1$ to match the growth overshoot) except to tune the surplus of $P$ by increasing $S_0$, which is in the opposite direction to tuning the surplus of $U$ [Fig. N6, Figs. S7 and S8].

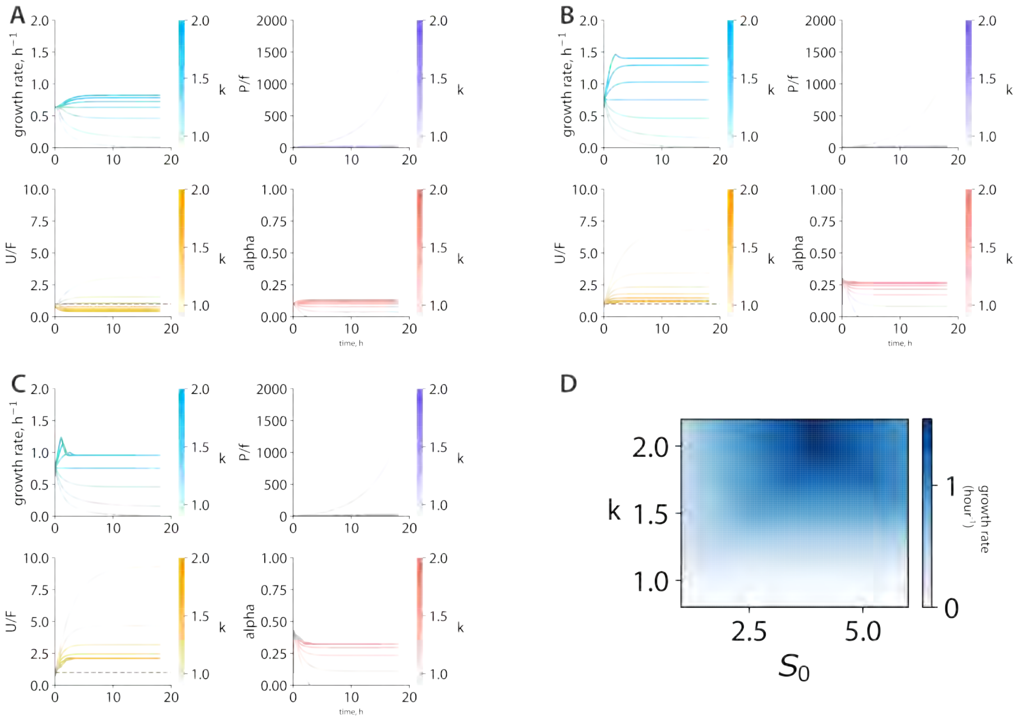

**Fig. N2: Numerical results of a steady-state condition. (A)-(C)** Different quantities, growth rate, $\frac{P}{f}$, $\frac{U}{F}$, and $\alpha$ change over time, by varying $k$ values when fixing $S_0$ at 2, 4.4, and 6, respectively. Other parameters are fixed as in Table SI. Note that the system is not at a steady state at the initial time point. **(D)** The steady-state growth rate as a function of $k$ and $S_0$.

##### D. Comparison with other bacterial growth models

For steady-state bacterial growth, our framework is essentially consistent with many other recent models, such as those in [1, 6–8]. Specifically, when assuming the catalyst (enzyme) is fewer than the substrate in each central biosynthetic process, for example, $U < F$ in Eq. S1, the corresponding equation reduces to $\frac{dU}{dt} = \frac{\alpha \cdot U}{\tau_U}$, which is similar to the formulation used for characterizing the production of ribosomal proteins in other models. Another noticeable difference of our model is the incorporation of nutrient quality. Unlike many other models that use an agglomerated metabolic rate to reflect on nutrient quality, our model directly depicts nutrient quality using the proportionality between nutrient molecule abundance and biomass, $k$, without altering the rate constants of intracellular biosynthetic processes (Eq. S4).

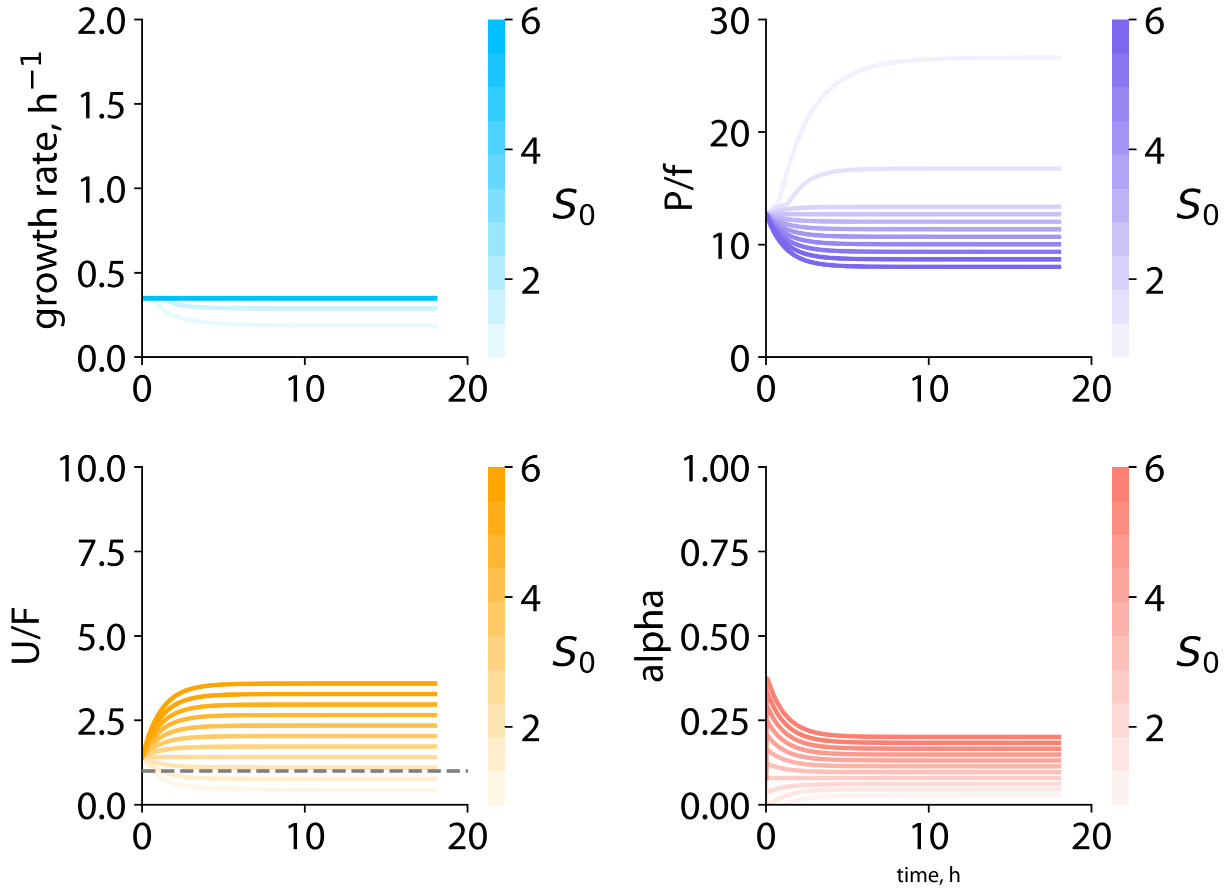

**Fig. N3: Numerical results of a steady-state condition.** Similar plots as in Fig. N2, by varying $S_0$ values when fixing $k$ at 1.2.

When characterizing the transitional dynamics between steady states, our model is also consistent with a recent work [5] that provides a mechanistic explanation of the oscillatory behaviors in growth adaptation during nutrient up-shift. In the Lagomarssino group's work, the key to reproducing the oscillatory dynamics is to introduce a mutual dynamic regulation between the the ribosome synthesis and the pool of the building blocks, i.e., amino acids, such as,

$$\frac{d\alpha}{dt} = \frac{1}{\tau_\alpha} \cdot \left[ \frac{[F]}{[F] + k_F} - \alpha(t) \right],$$
$$\frac{dF}{dt} = \frac{P}{\tau_F} - \frac{U}{\tau_U}, \tag{S7}$$

where the definitions of $\alpha$, $F$, $P$, $U$, $\tau_F$, and $\tau_U$ are the same as in Eqs. S1-S5, $[F]$ is the cellular concentration of $F$, $k_F$ is a Michaelis-constant equivalent and $\frac{1}{\tau_\alpha}$ is the rate of change

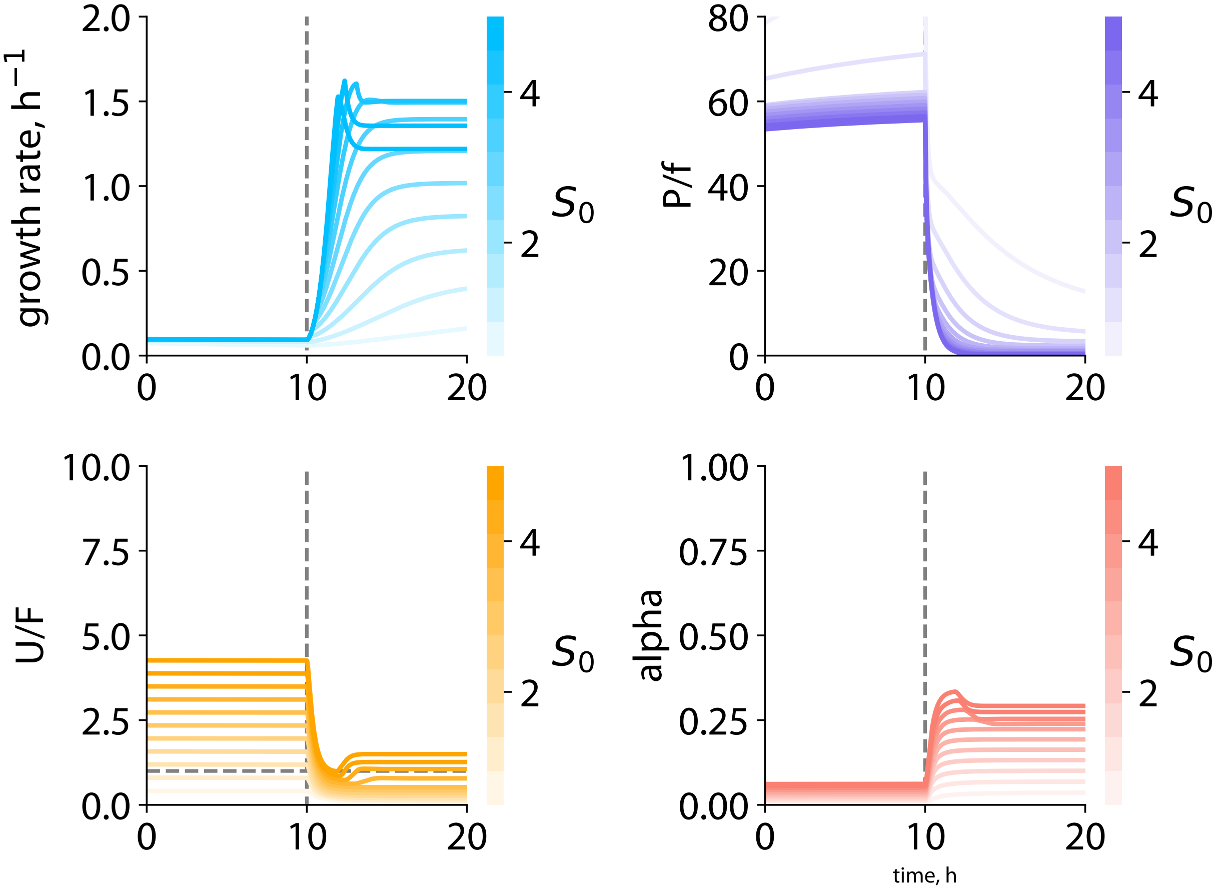

**Fig. N4: Numerical results of a nutrient up-shift to reproduce the *rpoC* as in Fig. 3B.** Similar plots as in Fig. N2, by shifting the $k$ value from 1.05 to 2.5 at the 10th hour, and by varying $S_0$ values to change the level of $U$. Decreasing $S_0$ corresponds to reducing $U$ and its surplus.

in the transcript pool, as defined in [5]. We used different annotations here from those in [5] to keep them consistent with our model description in Suppl. Notes I A. We also use absolute copy number terms here, such as $F$, $U$, and $P$, instead of the concentration used in the original work, again for consistency of formulation. Despite these differences, Eqs. S7 are mathematically equivalent to the formulations used in [5].

It is not difficult to show that Eqs. S7 have mathematical forms similar to those of Eqs.

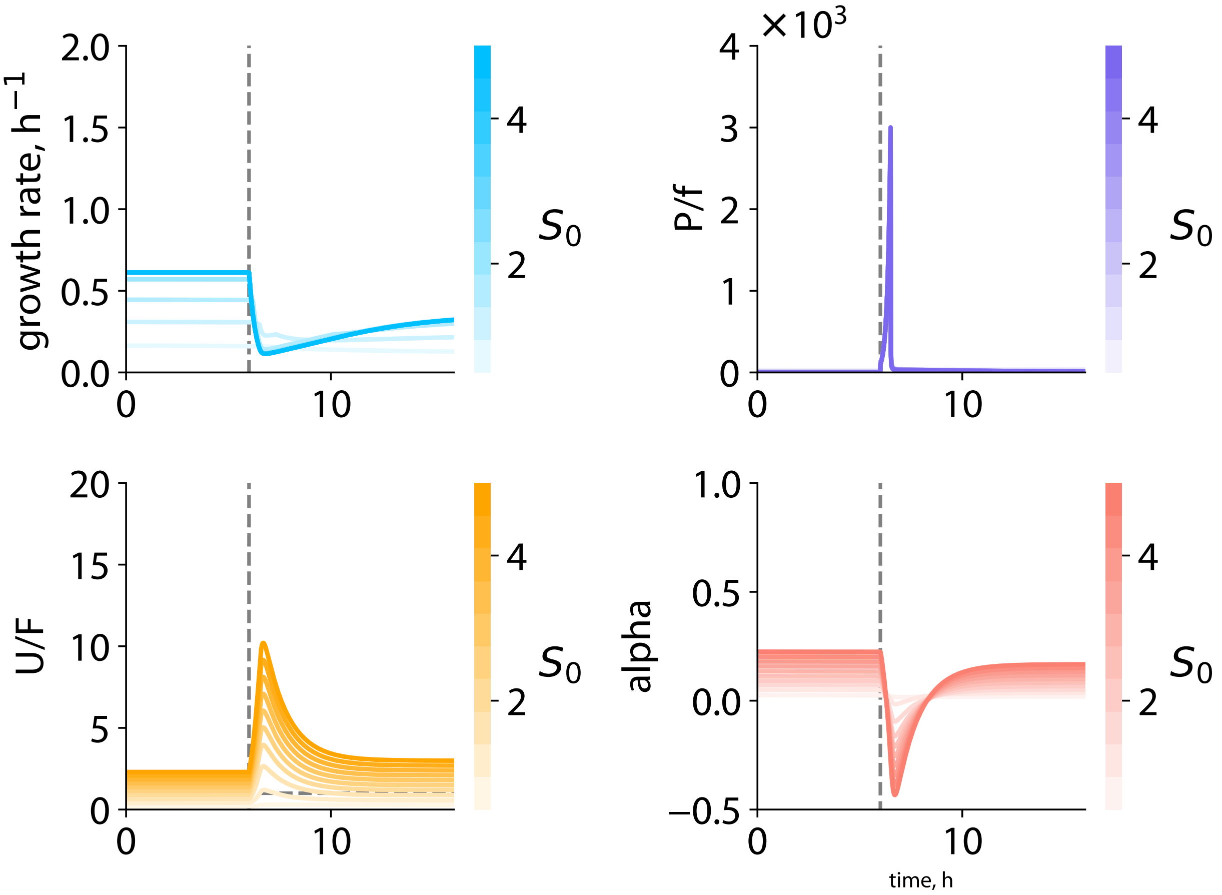

**Fig. N5: Numerical results of a nutrient down-shift to reproduce the *rpoC* as in Fig. 3C.** Similar plots as in Fig. N2, by shifting the $k$ value from 1.36 to 1.20 at the 6th hour, and by varying $S_0$ values to change the level of $U$. Decreasing $S_0$ corresponds to reducing $U$ and its surplus.

S1-S5, which are

$$\frac{d\alpha}{dt} = k_2 \left( S_0 - \frac{U}{F} \right) - k_1 \alpha,$$
$$\frac{dF}{dt} = \frac{\min(P, f)}{\tau_F} - \frac{\alpha \cdot \min(U, F)}{\tau_U} - \frac{(\phi_{\max} - \alpha) \cdot \min(U, F)}{\tau_P}. \tag{S8}$$

The similarity is more obvious after rearranging and assuming zero surplus for both $P$ and

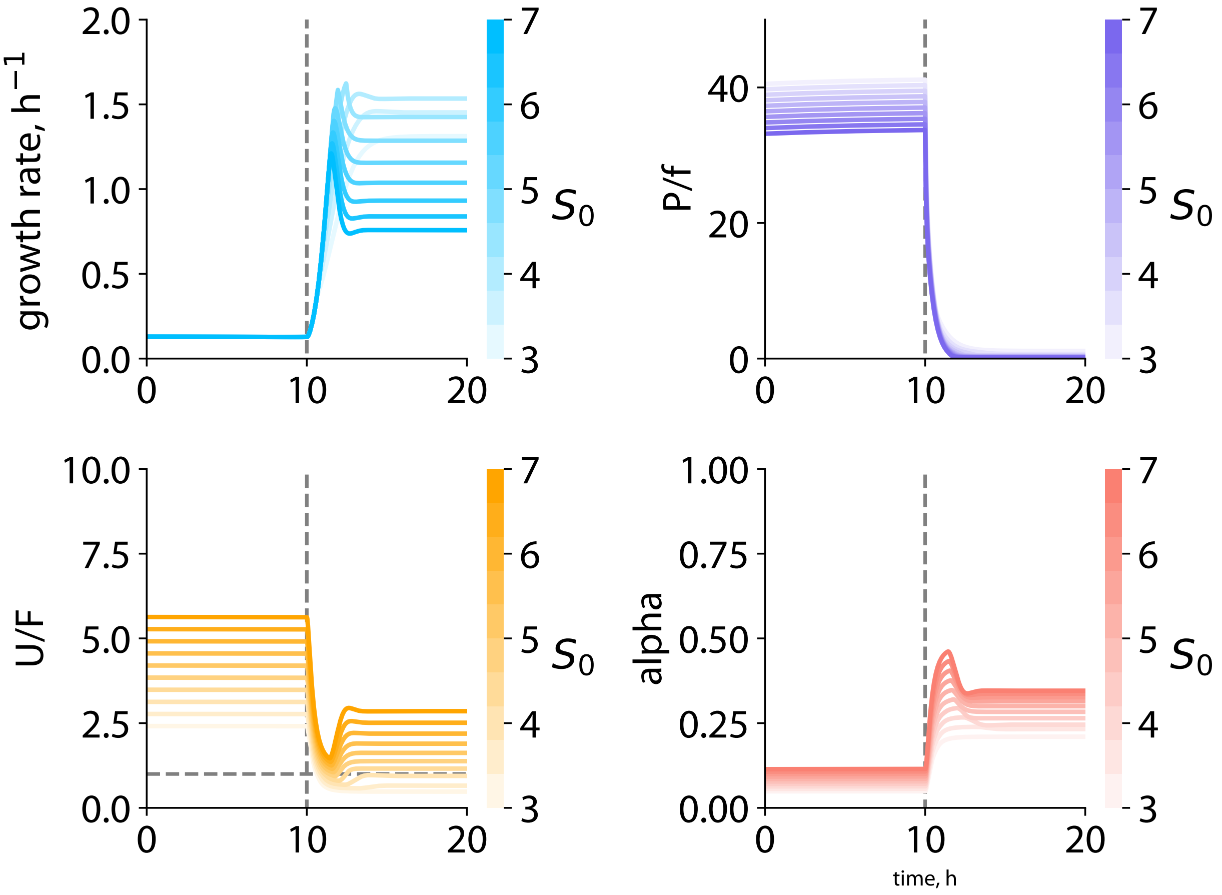

**Fig. N6: Numerical results of a nutrient up-shift to reproduce the *eno* as in Figs. S7 and S8.** Similar plots as in Fig. N2, by shifting the $k$ value from 1.08 to 2.5 at the 10th hour, and by varying $S_0$ values to change the level of $P$. Increasing $S_0$ corresponds to reducing $P$ and its surplus.

$U$, namely $P < f$ and $U < F$, such that,

$$\begin{aligned} \frac{d\alpha}{dt} &= k_2\left(S_0 - \frac{U}{F}\right) - k_1\alpha, \\ \frac{dF}{dt} &= \frac{P}{\tau_F} - \left(\frac{\alpha}{\tau_U} + \frac{\phi_{\max} - \alpha}{\tau_P}\right)\cdot U. \end{aligned} \tag{S9}$$

Both Eqs. S7 and S9 characterize the mutual regulatory scheme between $\alpha$ and $F$, specifically, $\alpha$ inhibiting $F$ while $F$ promoting $\alpha$. Within a certain range of rate constants $k_1$ and $k_2$ (gains), our model can reproduce the oscillatory behavior, despite different mathematical details, such as the specific dependence of $\frac{d\alpha}{dt}$ on $F$.

##### E. Depletion probability of core biosynthetic components within an autocatalytic network

Transcription and translation machines, as part of the core biosynthetic components, can be viewed as self-replicating agents within a larger autocalalytic network (see description in Suppl. Notes I A and [1]). In a recently established theory based on branching processes (see model details in [9]), self-replicating agents with finite lifetime and doubling time, both with stochasticity, have a non-zero probability of being completely depleted. The depletion probability, $P$, as a function of the number of agents, $N$, follows,

$$P(N) = \left(\frac{\Gamma_0}{\mu}\right)^{\alpha_r \cdot N}, \tag{S10}$$

where $\Gamma_0$ is the active degradation rate of the agents, inversely proportional to their lifetime, $\mu$ is the steady-state growth rate of the agents, and $\alpha_r$ is the fraction of the agents that process their own self-replication.

In the specific case of RNAP's self-replication, we can measure or estimate the values for all the parameters listed above. We applied Eq. S10 to the slowest steady-state growth condition where the surplus of RNAP is significant, and the active degradation rate for protein is non-negligible compared to the steady-state growth rate. Here, the measured growth rate $\mu$ in this slow growth condition is 0.16 hour$^{-1}$, the estimated number of RNAP, $N$, in that condition is about $2,000$ copies per cell [10, 11], the estimated active degradation rate, $\Gamma_0$, is about 0.083 hour$^{-1}$ [12, 13]. The only fitted parameter, the fraction of RNAP that transcribes *rpoB* or *rpoC*, $\alpha_r$, is about 0.8%, which matches well with the value measured by quantitative proteomics [10, 11]. Therefore, with only one free parameter, the calculated result by Eq. S10 is consistent with our experimental data (Fig. 4B).

Based on this model, we expect that the depletion probability of RNAP will be amplified when the fluctuation in the copy number of RNAP that transribes is own genes is larger. For examples, during nutrient up-shifts or growth-recovery after carbon starvation, the pool of free RNAPs faces stronger competition between binding to different sigma factors due to environmental shifts and therefore encounters larger fluctuations [REFs: Klumpp 2008, Yan 2024, TBA]. Indeed, our measured death probability, including the filamentation and cell lysis, is significantly increased in both cases [Fig. 4B, Fig. S10].

##### F. Justification of course-grained modeling of growth dynamics

There are several approaches for modeling how the cell allocates RNA-polymerase and ribosomes, from a detailed approach that attempts to describe in maximal details the actual mechanisms the cell employs for that purpose, to a coarse-grained approach that attempts to have a broad-stroke view of the process and the characterization and execution dilemmas faced by evolution from a bird's-eye view that is more general and universal but at a price that it neglects biological details of implementation. An example of the latter approach is the papers [14, 15] that demonstrated how product feedback inhibition is sufficient to lock the cell to optimal growth conditions, irrespective of the actual biological implementation, e.g., using riboswitches in tryptophan biosynthesis [16], or via the ppGpp mechanism in the stringent response [5, 17]. The details of the actual mechanism used by a particular cell often mask simple and universal design issues that are decoupled from the actual implementation. For example, a feedback mechanism that controls the growth rate by controlling the allocation of ribosomes toward self-replication, e.g., via product feedback inhibition, is expected to exhibit a single damped oscillation if it is optimally tuned to minimize the transition time irrespective of how the feedback is implemented. Furthermore, as we showed in previous sections, a feedback mechanism with low gain will cause the growth rate response to shift to be underdamped – slow in reaching the steady growth condition in the new environment. In contrast, a gain that is too high will cause excessive oscillations in the growth rate, which will relax to the steady growth condition also at a longer time scale compared to the optimum. Our simplified model enables these non-trivial predictions, which will not change even if more biological details are added to the model.

Furthermore, many cells have overlapping control mechanisms that perform the same task. This further complicates the detailed modeling approach. Here, we take the coarse-grained modeling approach. We wanted to understand to what extent we can capture the dynamic response to the upshift using a simple model. We found that our model can explain the observed behavior and, most importantly, shed light on the trade-off between the phenotypic adaptation rate and the steady-state growth rate.

#### II. SUPPLEMENTAL TABLES

| *E. coli* strains | Genotype | Experiments used | Notes |
|---|---|---|---|
| FS003 | K-12 MG1655 F- $\lambda$- rph-1 | – | background strain; low motility [18] |
| FS008 | FS003 with pLacIq::lacY A177C, pBAD_dCas9, ΔlacI, ΔaraE, ΔaraFGH::speC, galM//promoter tet sacB handle terminator//gmpA and pSIM18 | – | background strain for constructing CRISPRi cells |
| FS481 | FS003 ΔftsZ::[ftsZ-neongreen-Tn10] and pFS157 (pXY029-T5-pLAC mCherry-MinD MTSx2) | – | background strain for constructing *rpoD* rescue plasmid |
| HW050 | FS003 with pLacIq::lacY A177C, pBAD_dCas9, ΔlacI, ΔaraE, ΔaraFGH::speC, galM//promoter tet sacB handle terminator//gmpA, eno msfGFP cmR and pSIM18 | steady state growth in Figs. 2A-B, S2A, B and E, S4 and N9 | *eno* parental cell in this study |
| HW057 | FS003 with pLacIq::lacY A177C, pBAD_dCas9, ΔlacI, ΔaraE, ΔaraFGH::speC, galM//promoter enosgRNA handle terminator//gmpA, eno msfGFP cmR and pSIM18 | steady state growth and nutrient upshift experiments in Figs. 2A-B, S2A, B and E, S4, S7B-C, S8, S10B, N9 and N14, | *eno* CRISPRi cell in this study |

| | | | |
|---|---|---|---|
| HW002 | FS003 with pLacIq::lacY A177C, pBAD_dCas9, ΔlacI, ΔaraE, ΔaraFGH::speC, galM//promoter tet sacB handle terminator//gmpA, rpoB msfGFP cmR and pSIM18 | steady state growth in Figs. 2A-B, 4B-C, S2C, S4, N7 | *rpoB* parental cell in this study |
| HW006 | FS003 with pLacIq::lacY A177C, pBAD_dCas9, ΔlacI, ΔaraE, ΔaraFGH::speC, galM//promoter rpoBsgRNA handle terminator//gmpA, rpoB msfGFP cmR and pSIM18 | steady state growth, nutrient upshift and downshift experiments in Figs. 2A-B, 4B-C, S2C, S4, S6, S7C, S10A and N7 | *rpoB* CRISPRi cell in this study |
| HW102 | FS003 with pLacIq::lacY A177C, pBAD_dCas9, ΔlacI, ΔaraE, ΔaraFGH::speC, galM//promoter tet sacB handle terminator//gmpA, rpoC msfGFP cmR and pSIM18 | steady state growth in Figs. 1C-D, 2, 4B-C, S3A-C and S4 | *rpoC* parental cell with transcriptional reporter in this study |
| HW113 | FS003 with pLacIq::lacY A177C, pBAD_dCas9, ΔlacI, ΔaraE, ΔaraFGH::speC, galM//promoter rpoCsgRNA handle terminator//gmpA, rpoC msfGFP cmR and pSIM18 | steady state growth, nutrient upshift and downshift experiments in Figs. 1C-D, 2, 3B, 4B-C, S3A-C, S4, S5, S7C, S10A, N11 and N12 | *rpoC* CRISPRi cell with transcriptional reporter in this study |

| | | | |
|---|---|---|---|
| HW098 | FS003 with pLacIq::lacY A177C, pBAD_dCas9, ΔlacI, ΔaraE, ΔaraFGH::speC, galM//promoter tet sacB handle terminator//gmpA, rpoC-msfGFP cmR and pSIM18 | steady state growth in Figs. S2G and S3D | *rpoC* parental cell with translational reporter in this study |
| HW105 | FS003 with pLacIq::lacY A177C, pBAD_dCas9, ΔlacI, ΔaraE, ΔaraFGH::speC, galM//promoter rpoCsgRNA handle terminator//gmpA, rpoC-msfGFP cmR and pSIM18 | steady state growth, nutrient upshift and downshift experiments in Figs. S2G, S3D, N13 and N16 | *rpoC* CRISPRi cell with translational reporter in this study |
| HW010 | FS003 with pLacIq::lacY A177C, pBAD_dCas9, ΔlacI, ΔaraE, ΔaraFGH::speC, galM//promoter tet sacB handle terminator//gmpA, rpoH msfGFP cmR and pSIM18 | steady state growth in Figs. 2A-B, S2F, S4 and N10 | *rpoH* parental cell in this study |
| CA003 | FS003 with pLacIq::lacY A177C, pBAD_dCas9, ΔlacI, ΔaraE, ΔaraFGH::speC, galM//promoter rpoHsgRNA handle terminator//gmpA, rpoH msfGFP cmR and pSIM18 | steady state growth experiments in Figs. 2A-B, S2F, S4 and N10 | *rpoH* CRISPRi cell in this study |

| | | | |
|---|---|---|---|
| HW085 | FS003 with pLacIq::lacY A177C, pBAD_dCas9, ΔlacI, ΔaraE, ΔaraFGH::speC, galM//promoter tet sacB handle terminator//gmpA, tufA msfGFP cmR and pSIM18 | steady state growth in Figs. 2A-B, S2D, S4 and N8 | *tufA* parental cell in this study |
| HW091 | FS003 with pLacIq::lacY A177C, pBAD_dCas9, ΔlacI, ΔaraE, ΔaraFGH::speC, galM//promoter tufAsgRNA handle terminator//gmpA, tufA msfGFP cmR and pSIM18 | steady state growth, nutrient upshift and downshift experiments in Figs. 2A-B, S2D, S4, S7C, S10C, N8 and N15 | *tufA* CRISPRi cell in this study |

**Table SII: Strain Information of *E. coli.***

Note: '-' connected genes are fused with a linker, and will be co-transcribed in cells.

| Reported Gene | Media | Strain | Experimental parameters | Sample Size | Position in figures |
|---|---|---|---|---|---|
| *eno* | glucose + arginine | HW050 Parent | 0% Ara | 142218 | Figs. 2A-B, S2A, B and E, S4 and N9 |
| | | HW057 tCRISPRi | 0% Ara | 78116 | |
| | | | 0.005% Ara | 94962 | |
| | | | 0.05% Ara | 107189 | |
| | | | 0.1% Ara | 87350 | |
| | | | 0.2% Ara | 114827 | |
| | glucose | HW050 | 0% Ara | 78369 | |
| | | HW057 | 0% Ara | 76748 | |
| | | | 0.005% Ara | 161538 | |
| | | | 0.05% Ara | 116743 | |
| | | | 0.1% Ara | 52857 | |
| | | | 0.2% Ara | 35596 | |
| | rich glucose | HW050 | 0% Ara | 69787 | |
| | | HW057 | 0% Ara | 104218 | |
| | | | 0.005% Ara | 85308 | |
| | | | 0.05% Ara | 72518 | |
| *rpoB* | sodium acetate + arginine | HW002 | 0% Ara | 60473 | Figs. 2A-B, 4B-C, S2C, S4 and N7 |
| | | HW006 | 0% Ara | 27241 | |
| | | | 0.005% Ara | 1725 | |
| | | | 0.05% Ara | 18760 | |
| | | | 0.1% Ara | 7456 | |
| | | | 0.2% Ara | 4820 | |
| | glucose | HW002 | 0% Ara | 3406 | |
| | | HW006 | 0% Ara | 14902 | |

| | | | | | |
|---|---|---|---|---|---|
| | | | 0.005% Ara | 12681 | |
| | | | 0.05% Ara | 11002 | |
| | rich glucose | HW002 | 0% Ara | 58662 | |
| | | HW006 | 0% Ara | 87985 | |
| | | | 0.005% Ara | 67630 | |
| | | | 0.05% Ara | 87207 | |
| | | | 0.1% Ara | 62912 | |
| | | | 0.2% Ara | 50105 | |
| *rpoC* translational reporter | glucose | HW098 parent | 0% Ara | 82273 | Figs. S2G and S3D |
| | | HW105 tCRISPRi | 0% Ara | 87754 | |
| | | | 0.01% Ara | 40602 | |
| | | | 0.04% Ara | 21015 | |
| *rpoC* transcriptional reporter | sodium acetate + arginine | HW102 parent | 0% Ara | 120681 | Figs. 1C-D, 2, 4B-C, S3A-C and S4 |
| | | HW113 | 0% Ara | 15796 | |
| | | | 0.005% Ara | 23484 | |
| | | | 0.05% Ara | 13789 | |
| | | | 0.1% Ara | 11496 | |
| | | | 0.2% Ara | 17847 | |
| | glucose | HW102 parent | 0% Ara | 121784 | |
| | | HW113 | 0% Ara | 107679 | |
| | | | 0.005% Ara | 85054 | |
| | | | 0.05% Ara | 27069 | |
| | rich glucose | HW102 parent | 0% Ara | 47624 | |
| | | HW113 | 0% Ara | 86221 | |
| | | | 0.005% Ara | 77988 | |
| | | | 0.05% Ara | 35920 | |
| | | | 0.1% Ara | 17537 | |

| | | | | | |
|---|---|---|---|---|---|
| | | | 0.2% Ara | 2745 | |
| *rpoH* | glucose | HW010 parent | 0% Ara | 90583 | Figs. 2A-B, S2F, S4 and N10 |
| | | CA003 tCRISPRi | 0% Ara | 107867 | |
| | | | 0.005% Ara | 56115 | |
| | | | 0.01% Ara | 45209 | |
| *tufA* | sodium acetate + arginine | HW085 parent | 0% Ara | 76602 | Figs. 2A-B, S2D, S4 and N8 |
| | | HW091 tCRISPRi | 0% Ara | 44083 | |
| | | | 0.005% Ara | 20368 | |
| | | | 0.05% Ara | 24834 | |
| | | | 0.1% Ara | 30848 | |
| | | | 0.2% Ara | 43674 | |
| | glucose | HW085 parent | 0% Ara | 66299 | |
| | | HW091 tCRISPRi | 0% Ara | 59717 | |
| | | | 0.005% Ara | 114846 | |
| | | | 0.05% Ara | 75831 | |
| | rich glucose | HW085 parent | 0% Ara | 14498 | |
| | | HW091 tCRISPRi | 0% Ara | 95955 | |
| | | | 0.005% Ara | 78546 | |
| | | | 0.05% Ara | 121793 | |
| | | | 0.1% Ara | 95467 | |
| | | | 0.2% Ara | 44145 | |

**Table SIII: Steady state experimental conditions and sample size. Related to the STAR Methods.**

Each main trench on the mother machine microfluidics contains 4000 individual channels. After cells were injected into the main trench, cells were loaded into channels by centrifuging the device. The sample size represents the number of transient single-cell growth properties measured from each mother machine experiment. '$\rightarrow$' indicates that the shift experiment was conducted between the two concentrations of inducer on both sides.

| Reported Gene | Media | Strain | Experimental parameters | Sample Size | Position in figures |
|---|---|---|---|---|---|
| *eno* | glucose + arginine → rich glucose | HW057 tCRISPRi | 0% Ara | 49963 | Figs. S7B-C and S8 |
| | | | 0.01% Ara | 101298 | |
| | | | 0.04% Ara | 114629 | |
| | | | 0.08% Ara | 80787 | |
| | | | 0.04% Ara + pyruvate | 133672 | |
| | mannose → rich glucose | HW057 tCRISPRi | 0% Ara | 58233 | Fig. N14 |
| | | | 0.01% Ara | 40713 | |
| | | | 0.02% Ara → 0% Ara | 49081 | |
| *rpoB* | sodium acetate + arginine → rich glucose | HW006 tCRISPRi | 0% Ara | 3109 | Figs. S6, S7C and S10A |
| | | | 0.04% Ara | 42331 | |
| *rpoC* translational reporter | sodium acetate + arginine → rich glucose | HW105 tCRISPRi | 0% Ara | 1864 | Fig. N13 |
| | | | 0.04% Ara | 10832 | |
| *rpoC* transcriptional reporter | sodium acetate + arginine → rich glucose | HW113 tCRISPRi | 0% Ara | 16714 | Figs. 3B, S5, S7C and S10A |
| | | | 0.04% Ara | 5956 | |
| | sodium acetate + arginine → rich glucose | HW113 tCRISPRi | 0% Ara | 41338 | Fig. N11 |
| | | | 0.04% Ara | 18220 | |
| | | | 0.04% Ara → 0% Ara | 12043 | |
| | | | 0.04% Ara + *rpoD* | 15607 | |
| | sodium acetate → rich glucose | HW113 tCRISPRi | 0% Ara | 31275 | Fig. N12 |
| | | | 0.04% Ara | 5097 | |
| *tufA* | sodium acetate + arginine → rich glucose | HW091 tCRISPRi | 0% Ara | 83087 | Figs. S7C, S10C and N15 |
| | | | 0.04% Ara | 49084 | |

**Table S**IV: **Medium upshift experimental conditions and sample size. Related to the STAR Methods.**

The sample size represents the number of transient single-cell growth properties measured from each mother machine experiment. '$\rightarrow$' indicates that the shift experiment was conducted between the two concentrations of inducer on both sides.

| Reported Gene | Media | Strain | Experimental parameters | Sample Size | Position in main figures |
|---|---|---|---|---|---|
| *rpoC* translational reporter | glucose + 11 a.a. → sodium acetate | HW105 tCRISPRi | 0% Ara<br>0.01% Ara | 59742<br>28752 | Fig. N16 |
| *rpoC* translational reporter | glucose + 11 a.a. → sodium acetate | HW105 tCRISPRi | 0% Ara<br>0.05% Ara | 77752<br>43144 | Fig. N16 |
| *rpoC* transcriptional reporter | glucose + 11 a.a. → sodium acetate | HW113 tCRISPRi | 0% Ara<br>0.005% Ara | 101369<br>28544 | Fig. S9 |
| *rpoC* transcriptional reporter | glucose + 11 a.a. → sodium acetate | HW113 tCRISPRi | 0% Ara<br>0.05% Ara | 33411<br>12607 | Figs. 3C and S9 |

**Table SV: Medium downshift experimental conditions and sample size. Related to the STAR Methods.**

The symbols in the rightmost columns are the same as those in the corresponding main figures. '→' indicates that the shift experiment was conducted between the two concentrations of inducer on both sides.

| Media name (as used in the text) | Buffer | Carbon source (v/w) concentration | Supplement |
|---|---|---|---|
| glucose | MOPS modified buffer | Glucose 0.2% | – |
| glucose +11 a.a. | MOPS modified buffer | Glucose 0.2% | table SIX |
| rich glucose | MOPS modified buffer | Glucose 0.2% | table SVIII |
| sodium acetate | MOPS modified buffer | Sodium acetate 60 mM | – |
| sodium acetate + arginine | MOPS modified buffer | Sodium acetate 60 mM | No NH4Cl<br>L-arginine (R) 500 $\mu$g/ml |

**Table** SVI: **List of growth media, carbon scources and the supplements that we used for this study.**

| Components | Concentration |
|---|---|
| MOPS (MW 209.3) | 40mM |
| Tricine (MW 179.2) | 4.0 mM |
| Iron(II) Sulfate Stock | 0.1 mM |
| Ammonium Chloride | 9.5 mM |
| Sodium Sulfate | 0.276 mM |
| Calcium Chloride | 0.0005 mM |
| Magnesium Chloride | 0.525 mM |
| Sodium Chloride | 50mM |
| Ammonium Molybdate | $3\times10^{-9}$ M |
| Boric Acid | $4\times10^{-7}$ M |
| Cobalt Chloride | $3\times10^{-8}$ M |
| Cupric Sulfate | $10^{-8}$ M |
| Manganese Chloride | $8\times10^{-8}$ M |
| Zinc Sulfate | $10^{-9}$ M |
| Potassium Phosphate Monobasic | 1.32 mM |

**Table SVII: Components of MOPS modified buffer**

| Components | Concentration |
| --- | --- |
| L-Alanine | 0.8 mM |
| L-Arginine | 5.2 mM |
| L-Asparigine | 0.4 mM |
| L-Aspartic Acid, Potassium Salt | 0.4 mM |
| L-Glutamic Acid, Potassium Salt | 0.66 mM |
| L-Glutamine | 0.6 mM |
| L-Glycine | 0.8 mM |
| L-Histidine HCl $H_2O$ | 0.2 mM |
| L-Isoleucine | 0.4 mM |
| L-Proline | 0.4 mM |
| L-Serine | 10 mM |
| L-Threonine | 0.4 mM |
| L-Tryptophan | 0.1 mM |
| L-Valine | 0.6 mM |
| L-Leucine | 0.8 mM |
| L-Lysine | 0.4 mM |
| L-Methionine | 0.2 mM |
| L-Phenylalinine | 0.4 mM |
| L-Cysteine HCl | 0.1 mM |
| L-Tyrosine | 0.2 mM |
| Thiamine | 0.01 mM |
| Calcium Pantothenate | 0.01 mM |
| para-Amino Benzoic Acid | 0.01 mM |
| para-Hydroxy benzoic Acid | 0.01 mM |
| di Hydroxy Benzoic Acid | 0.01 mM |
| Potassium Hydroxide | 1.5 mM |
| Adenine | 0.2 mM |
| Cytosine | 0.2 mM |
| Uracil | 0.2 mM |
| Guanine | 0.2 mM |

**Table SVIII: Supplements for synthetic rich media.**

| Components | Concentration ($\mu$g/ml) |
|---|---|
| L-methionine (M) | 500 |
| L-histidine (H) | 500 |
| L-arginine (R) | 500 |
| L-proline (P) | 500 |
| L-threonine (T) | 500 |
| L-tryptophan (W) | 500 |
| L-leucine (L) | 500 |
| L-tyrosine (Y) | 500 |
| L-alanine (A) | 500 |
| L-asparagine (N) | 500 |
| L-aspartic acid (D) | 25 |

**Table S**IX: Supplements for Glucose + 11 a.a.

#### III. SUPPLEMENTAL FIGURES

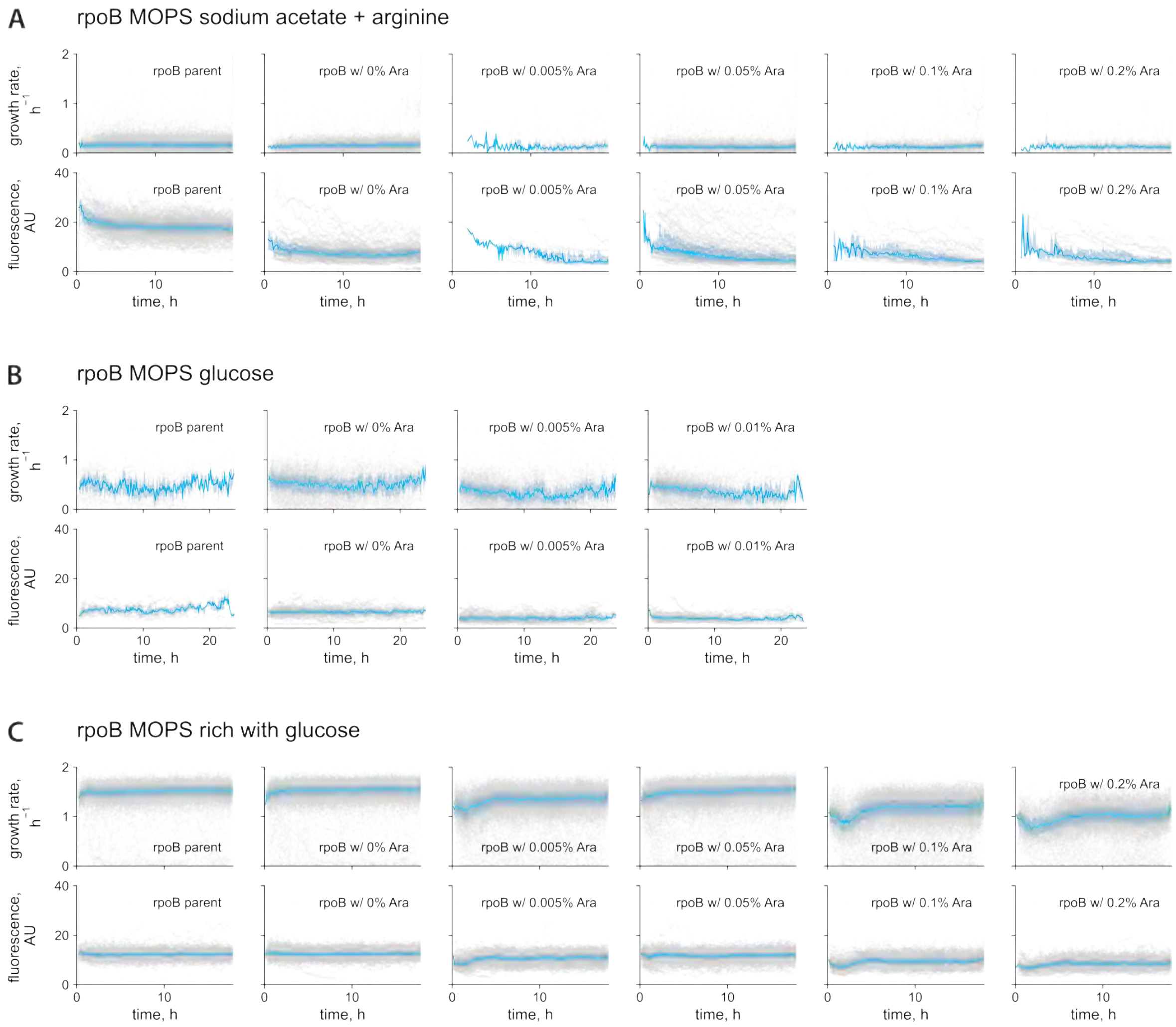

**Fig. N7: Growth-rate time course and fluorescent-intensity time course for steady-state growth of *rpoB* cells with transcriptional reporter in different growth media.** The scatter plot (light gray) shows single-cell growth rate versus time. The blue line represents the median growth rate, while the shaded blue area indicates the interquartile range. The same plotting style applies to Figs. N8-N10.

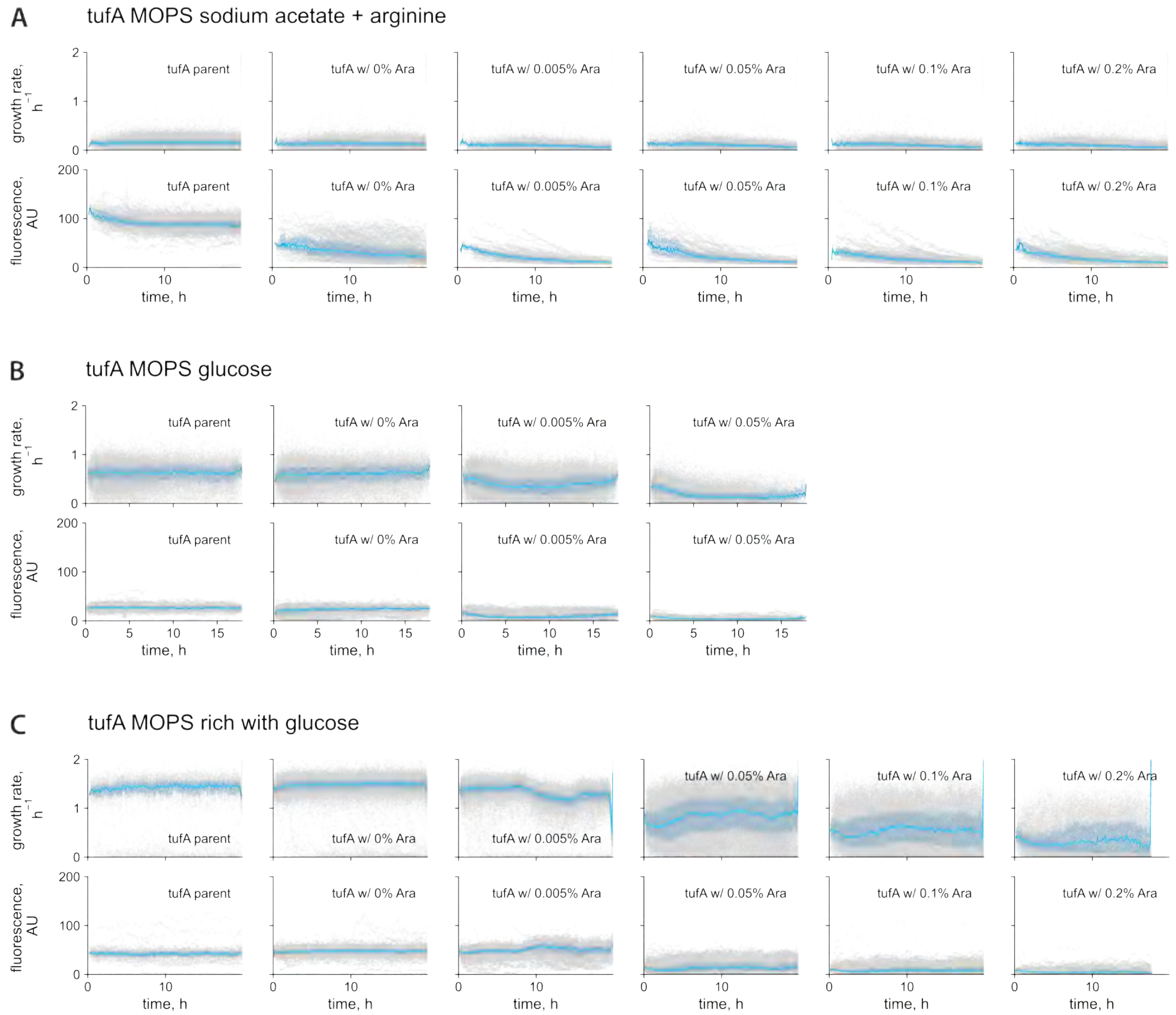

**Fig. N8: Growth-rate time course and fluorescent-intensity time course for steady-state growth of *tufA* cells with transcriptional reporter in different growth media.**

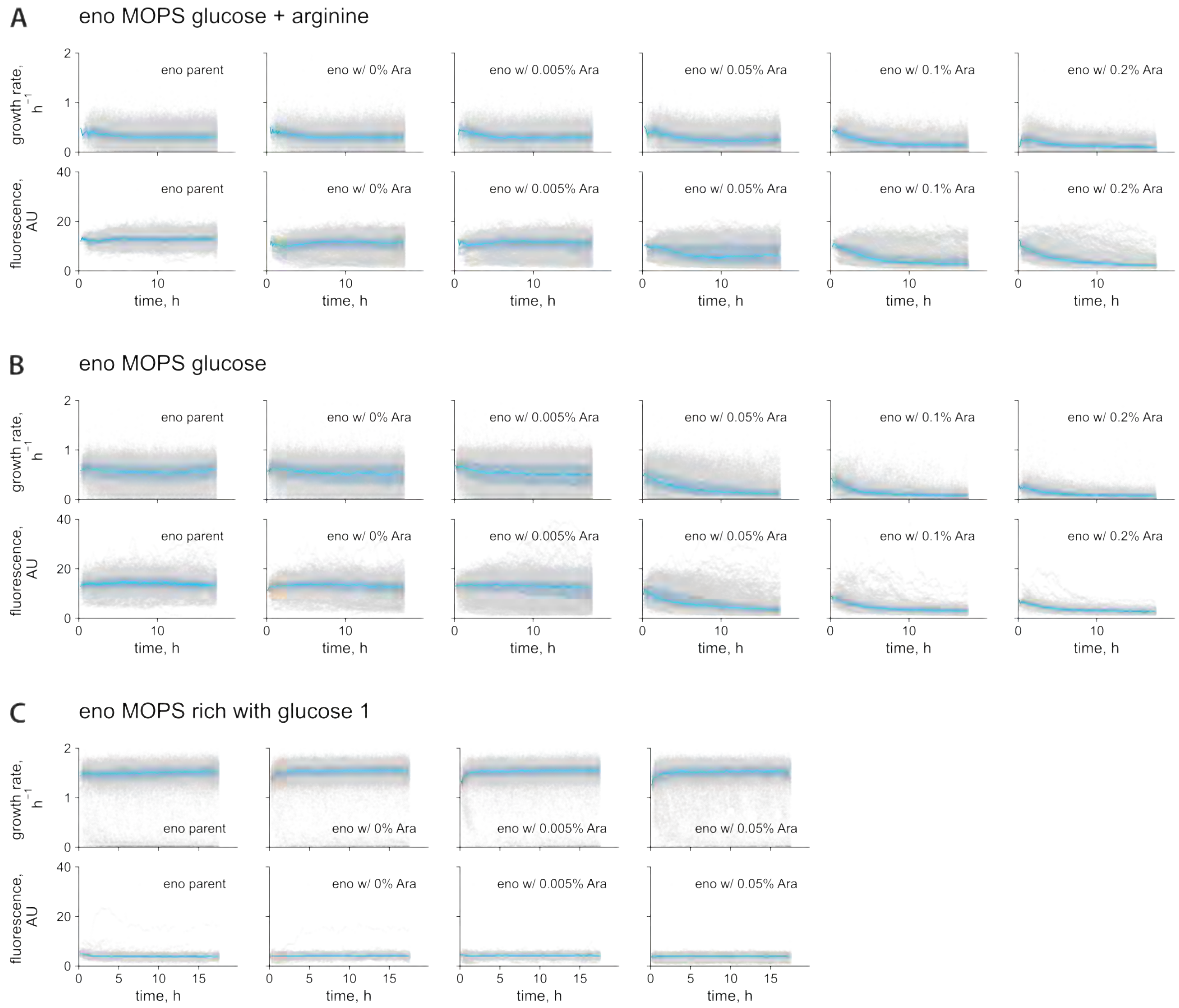

**Fig. N9: Growth-rate time course and fluorescent-intensity time course for steady-state growth of *eno* cells with transcriptional reporter in different growth media.**

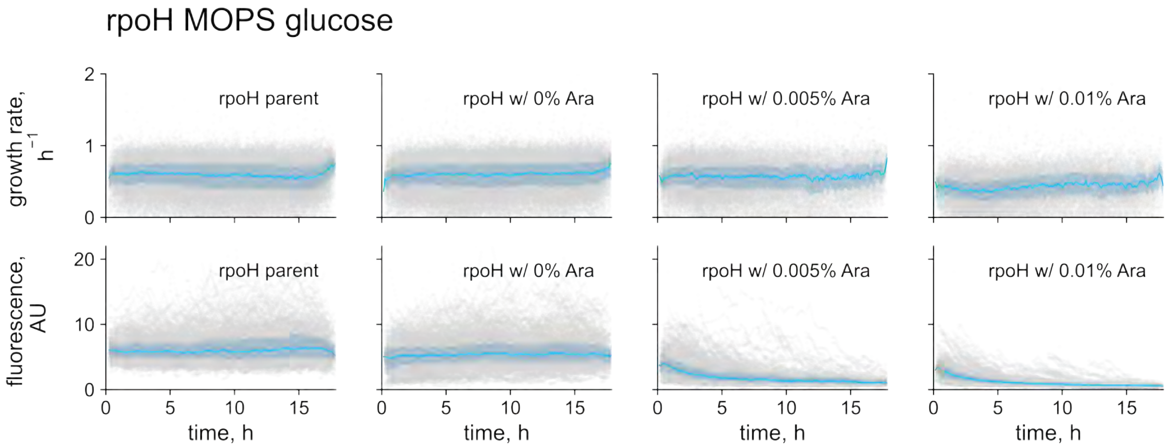

**Fig. N**10: **Growth-rate time course and fluorescent-intensity time course for steady-state growth of *rpoH* cells with transcriptional reporter in different growth media.**

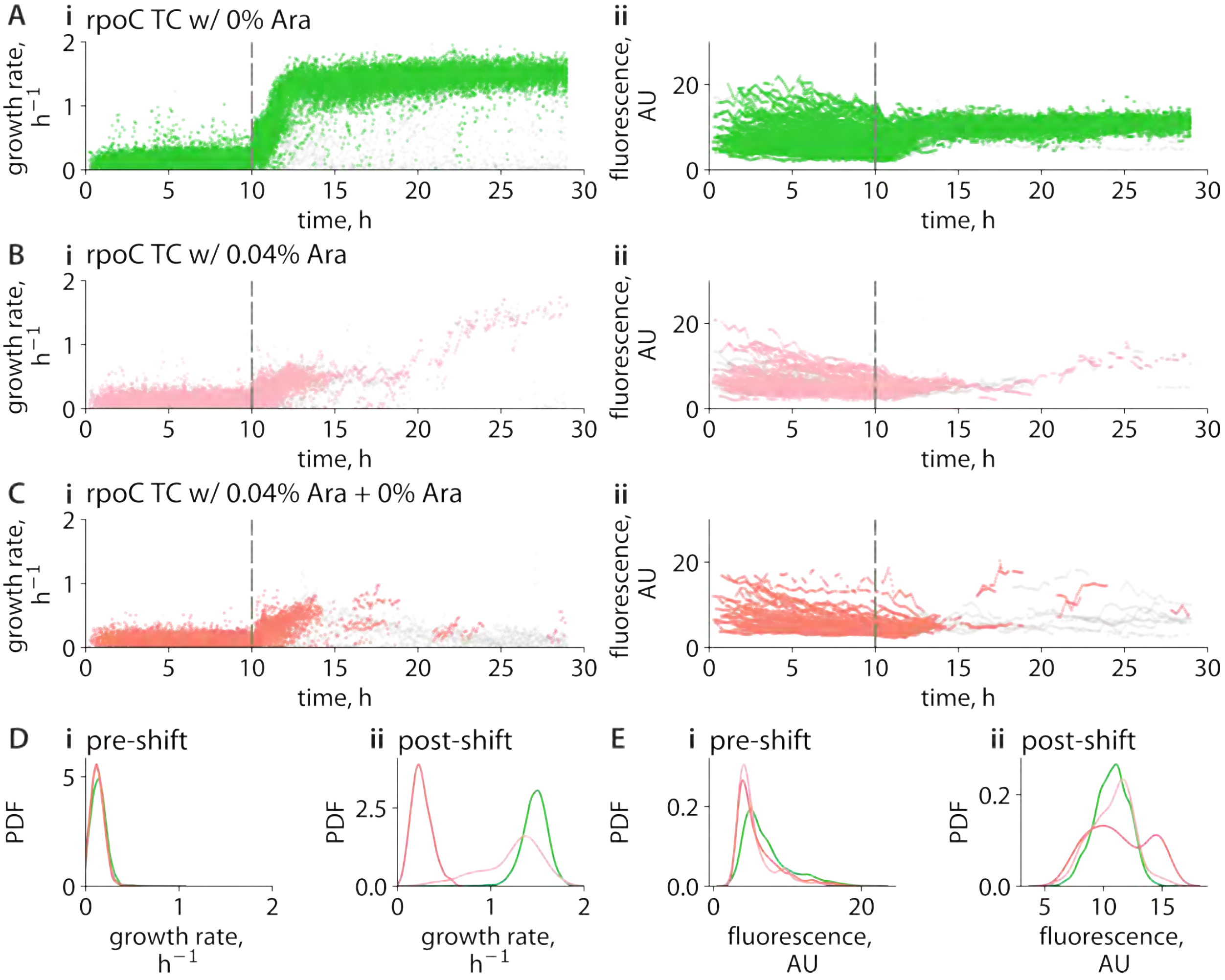

**Fig. N**11: Nutrient up-shift responses of *rpoC* cells when RpoC surplus is repressed in various conditions. In the up-shift experiments, cells were initially grown for 10 h at steady state in a slow-growth medium (MOPS supplemented with sodium acetate and arginine), followed by an immediate switch to a fast-growth medium (MOPS rich with glucose). **A–C,** time-course profiles of (i) single-cell growth rate (h$^{-1}$) and (ii) fluorescence intensity (AU) for *rpoC* cells under different conditions: **A,** no arabinose (Ara, 0%); **B,** 0.04% Ara; **C,** 0.04% Ara in the pre-shift condition and 0% Ara in the post-shift condition. Colored scatter points represent adapting cells; gray points indicate non-adapting cells. **D,** PDFs of single-cell growth rates before (i) and after (ii) the nutrient shift. **E,** PDFs of single-cell fluorescence intensities before (i) and after (ii) the shift. Color codes are consistent with panels A–C.

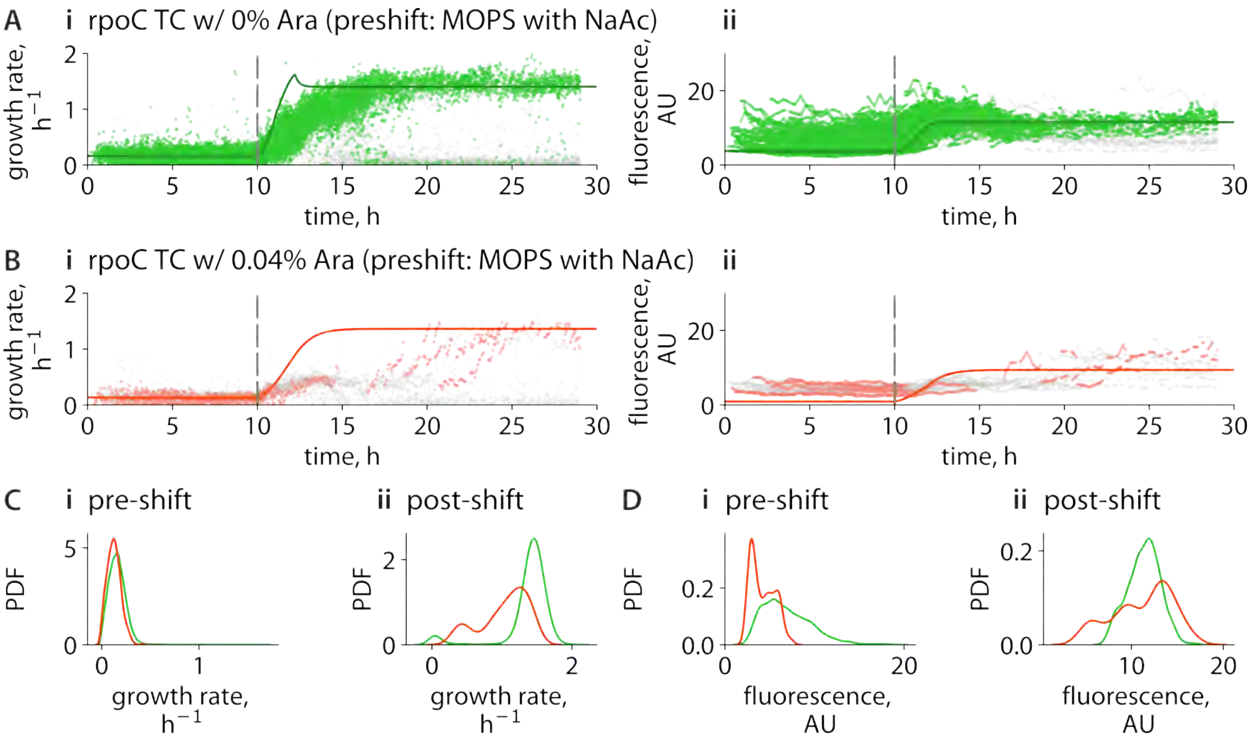

**Fig. N12: Nutrient up-shift experiments with *rpoC* cells show delays in adaptation when RpoC surplus is repressed in an alternative nutrient-poor medium.** In the up-shift experiments, cells were initially grown for 10 hours at steady state in a slow-growth medium (MOPS supplemented with sodium acetate, NaAc), followed by an immediate switch to a fast-growth medium (MOPS rich with glucose). **A.** Time-course profiles of (i) growth rate (hour$^{-1}$) and (ii) fluorescence intensity in arbitrary units (AU) for *rpoC* (transcriptional reporter, TC) cells cultured without arabinose (Ara) are shown. The green scatters represent the single-cell data for growing cells. Cells that did not adapt to the enriched medium are shown in gray. Dark green lines represent fitted dynamics using our model. Model details and parameters can be found in Supplemental Information and Table S1. **B.** Time-course profiles of (i) growth rate (hour$^{-1}$) and (ii) fluorescence intensity in AU for cells cultured with 0.04% arabinose. Red scatters indicate adapting cells; gray scatters represent non-adapting cells. Dark red lines show model fits. Plot styles are the same as A. **C.** Probability density distribution (PDF) of single-cell growth rate for pre-shift and post-shift conditions, estimated using Gaussian kernel density estimation, with green line for 0% Ara and red for 0.04% Ara. **D.** PDF of single-cell fluorescence intensities for pre-shift and post-shift conditions. Plot styles are the same as C.

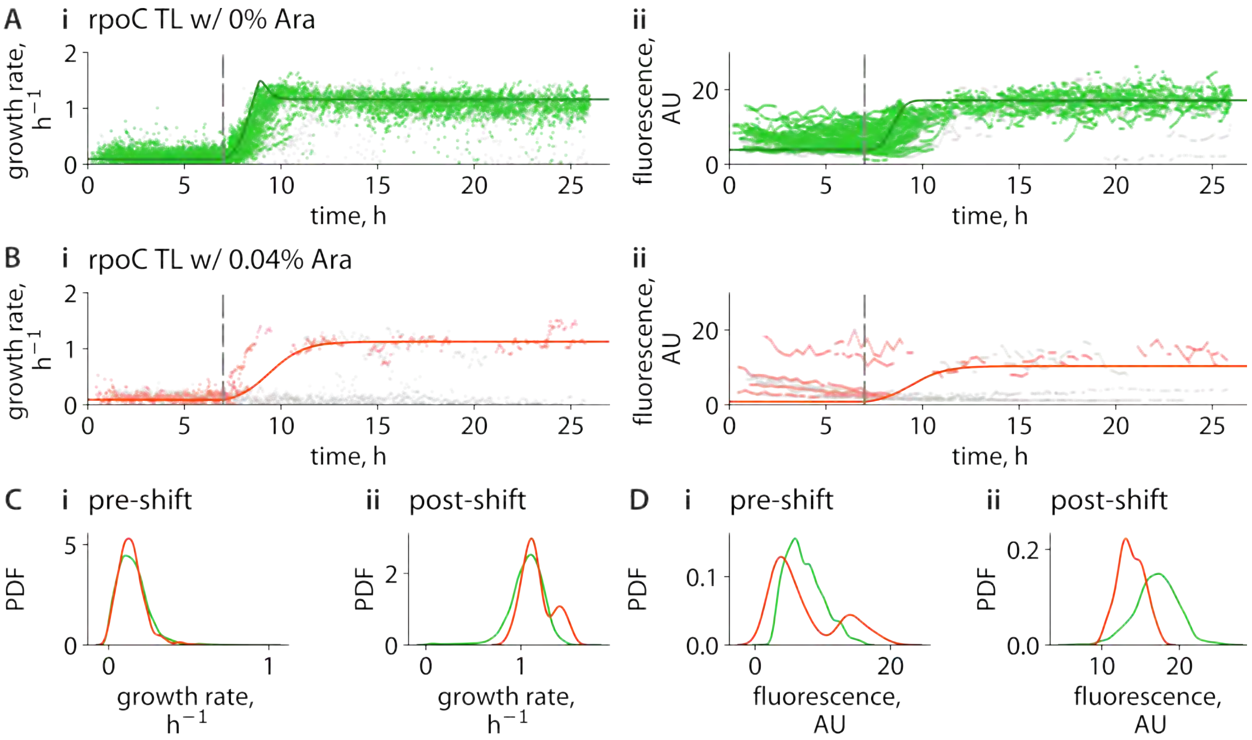

**Fig. N13: Nutrient up-shift experiments with *rpoC* cells with a translational reporter show delays in adaptation when RpoC surplus is repressed.** In the up-shift experiments, cells were initially grown for 10 hours at steady state in a slow-growth medium (MOPS supplemented with sodium acetate and arginine), followed by an immediate switch to a fast-growth medium (MOPS rich with glucose). **A.** Time-course profiles of (i) growth rate (hour$^{-1}$) and (ii) fluorescence intensity in arbitrary units (AU) for *rpoC* (translational reporter, TL) cells cultured without arabinose (Ara) are shown. The green scatters represent the single-cell data for growing cells. Cells that did not adapt to the enriched medium are shown in gray. Dark green lines represent fitted dynamics using our model. Model details and parameters can be found in Supplemental Information and Table S1. **B.** Time-course profiles of (i) growth rate (hour$^{-1}$) and (ii) fluorescence intensity in AU for cells cultured with 0.04% arabinose. Red scatters indicate adapting cells; gray scatters represent non-adapting cells. Dark red lines show model fits. Plot styles are the same as A. **C.** Probability density distribution (PDF) of single-cell growth rate for pre-shift and post-shift conditions, estimated using Gaussian kernel density estimation, with green line for 0% Ara and red for 0.04% Ara. **D.** PDF of single-cell fluorescence intensities for pre-shift and post-shift conditions. Plot styles are the same as C.

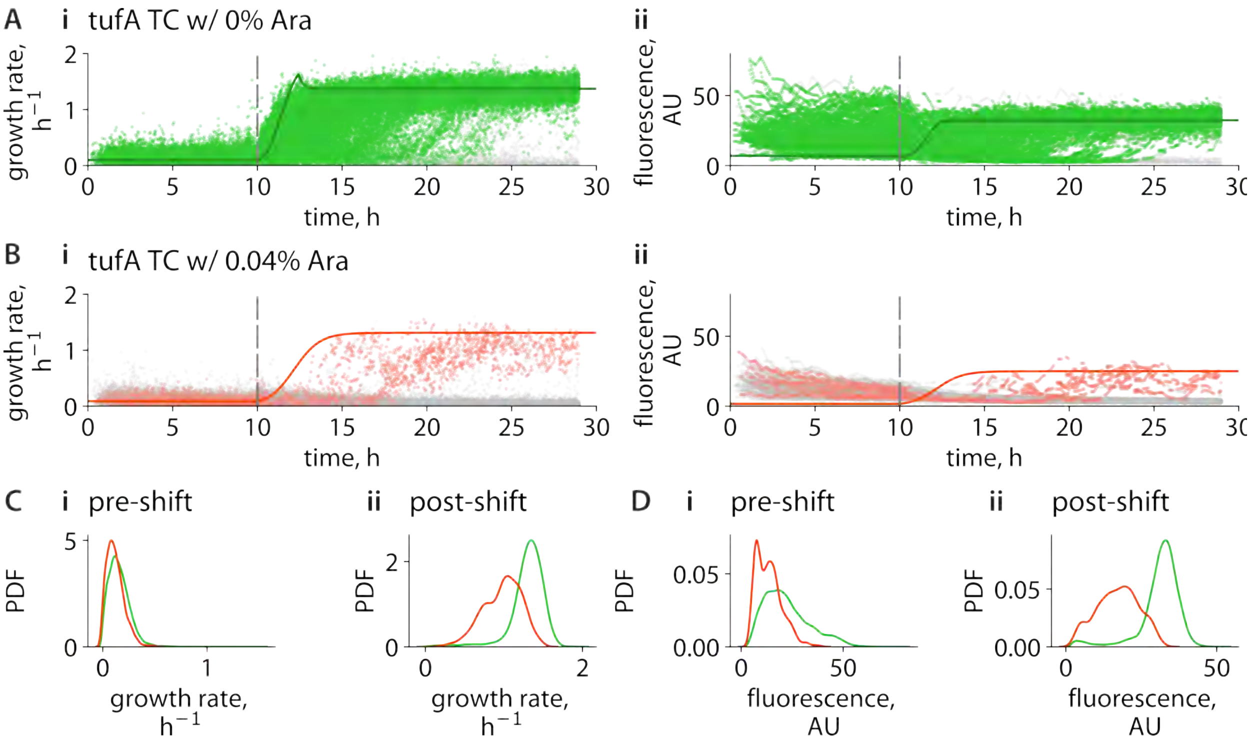

**Fig. N14: Nutrient up-shift experiments with *tufA* cells show delays in adaptation when TufA surplus is repressed.** In the up-shift experiments, cells were initially grown for 10 hours at steady state in a slow-growth medium (MOPS supplemented with sodium acetate and arginine), followed by an immediate switch to a fast-growth medium (MOPS rich with glucose). **A.** Time-course profiles of (i) growth rate ($\text{hour}^{-1}$) and (ii) fluorescence intensity in arbitrary units (AU) for *tufA* (transcriptional reporter, TC) cells cultured without arabinose (Ara) are shown. The green scatters represent the single-cell data for growing cells. Cells that did not adapt to the enriched medium are shown in gray. Dark green lines represent fitted dynamics using our model. Model details and parameters can be found in Supplemental Information and Table S1. **B.** Time-course profiles of (i) growth rate ($\text{hour}^{-1}$) and (ii) fluorescence intensity in AU for cells cultured with 0.04% arabinose. Red scatters indicate adapting cells; gray scatters represent non-adapting cells. Dark red lines show model fits. Plot styles are the same as A. **C.** Probability density distribution (PDF) of single-cell growth rate for pre-shift and post-shift conditions, estimated using Gaussian kernel density estimation, with green line for 0% Ara and red for 0.04% Ara. **D.** PDF of single-cell fluorescence intensities for pre-shift and post-shift conditions. Plot styles are the same as C.

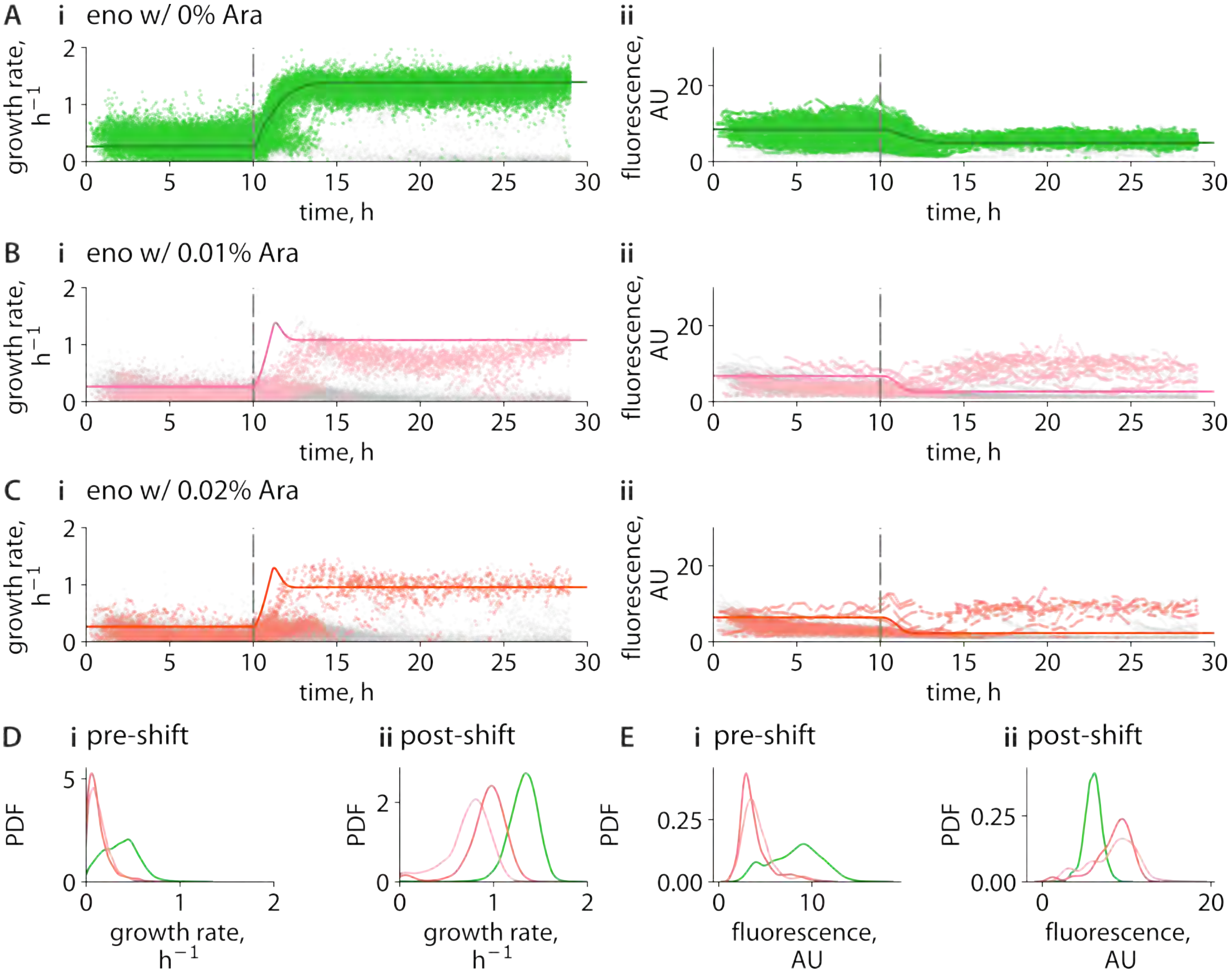

**Fig. N15: Nutrient up-shift experiments reveal invariant adaptation upon repression of *eno* surplus expression.** Cells were grown to steady state for 10 h in a slow-growth medium (MOPS supplemented with mannose) and then immediately transferred to a rich-growth medium (MOPS rich with glucose). **A–C,** Time courses of single-cell growth rate ($h^{-1}$; left) and *eno* translational reporter fluorescence (AU; right) following the nutrient up-shift in the presence of 0%, 0.01%, and 0.02% arabinose (Ara), respectively. Colored points denote adapting cells, gray points denote non-adapting cells, and solid lines show fits of the quantitative model (Supplementary Information and Table S1). **D,** Probability density distributions of single-cell growth rates before and after the shift, estimated by Gaussian kernel density estimation. **E,** Probability density distributions of single-cell fluorescence intensities before and after the shift. In **D** and **E**, green, pink, and red correspond to 0%, 0.01%, and 0.02% Ara, respectively.

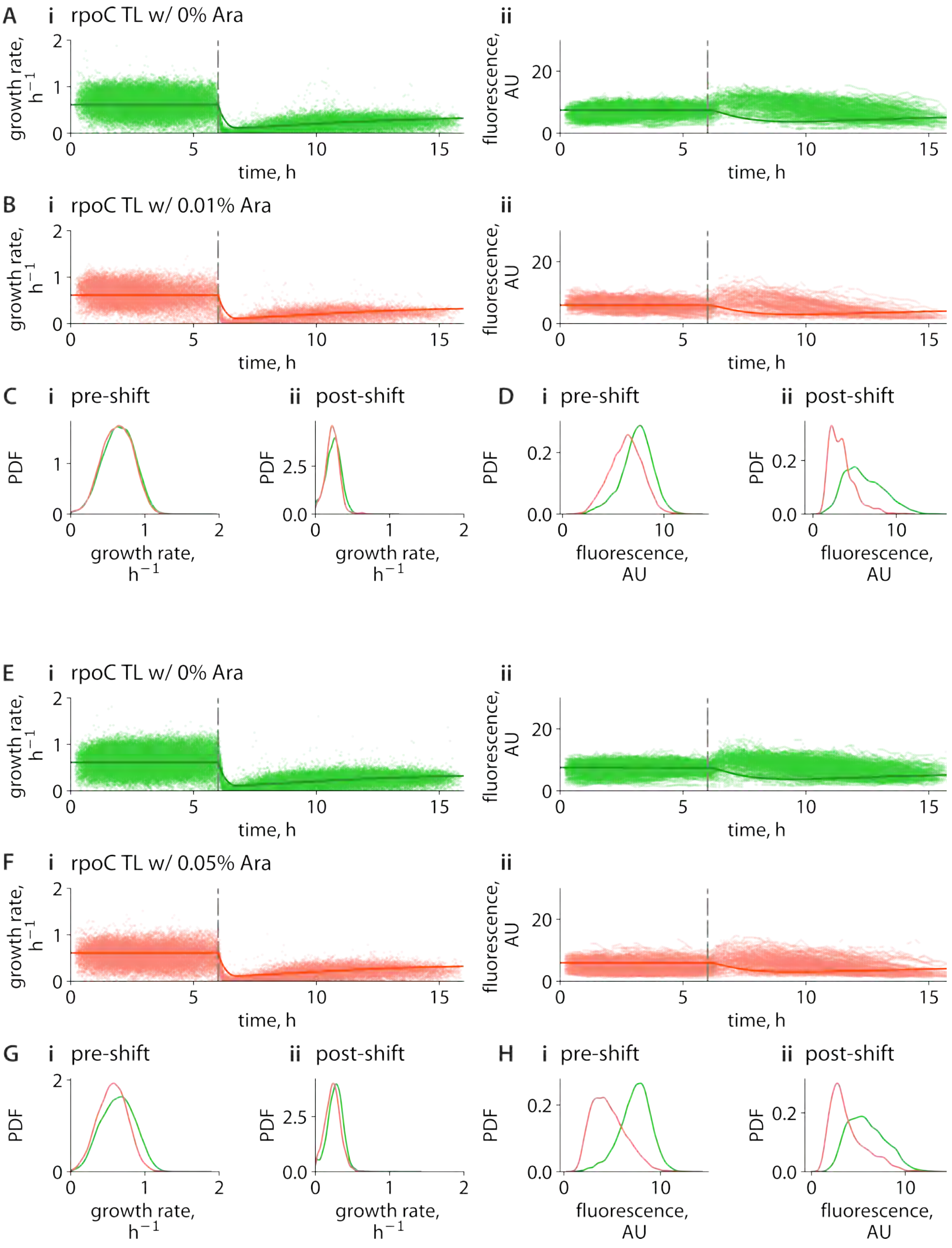

**Fig. N16: Nutrient down-shift experiments with *rpoC* translational reporter (TL) cells when RpoC surplus is repressed.**

**Fig. N16 (continued).** In the down-shift experiments, cells were initially grown for 6 hours at steady state in a fast-growth medium (MOPS supplemented with 11 amino acids and glucose), followed by an immediate switch to a slow-growth medium (MOPS with sodium acetate). A–D and E–H are two sets of experiments. **A–B.** Time-course profiles of (i) growth rate (hour$^{-1}$) and (ii) fluorescence intensity in arbitrary units (AU) for *rpoC* cells. **A.** Down-shift experiment time course with no arabinose. **B.** Down-shift experiment time course with 0.01% Ara. Plot styles are the same as A. **C.** PDFs of single-cell growth rates before (i) and after (ii) the nutrient shift, with green line for 0% Ara and red for 0.01% Ara. **D.** PDFs of single-cell fluorescence intensities before (i) and after (ii) the shift. **E–F.** Time-course profiles of (i) growth rate (hour$^{-1}$) and (ii) fluorescence intensity in arbitrary units (AU) for *rpoC* cells. **E.** Down-shift experiment time course with no arabinose. **F.** Down-shift experiment time course with 0.05% Ara. Plot styles are the same as E. **G.** PDFs of single-cell growth rates before (i) and after (ii) the nutrient shift, with green line for 0% Ara and red for 0.05% Ara. **H.** PDFs of single-cell fluorescence intensities before (i) and after (ii) the shift.

[1] A. Roy, D. Goberman, and R. Pugatch, A unifying autocatalytic network-based framework for bacterial growth laws, Proceedings of the National Academy of Sciences of the United States of America **118**, e2107829118 (2021).

[2] M. Scott, C. W. Gunderson, E. M. Mateescu, Z. Zhang, and T. Hwa, Interdependence of Cell Growth and Gene Expression: Origins and Consequences, Science **330**, 1099 (2010).

[3] X. Dai, M. Zhu, M. Warren, R. Balakrishnan, V. Patsalo, H. Okano, J. R. Williamson, K. Fredrick, Y.-P. Wang, and T. Hwa, Reduction of translating ribosomes enables Escherichia coli to maintain elongation rates during slow growth, Nature Microbiology **2**, nmicrobiol2016231 (2016).

[4] H. Bremer and P. P. Dennis, Modulation of chemical composition and other parameters of the cell by growth rate, (1996).

[5] R. Droghetti, P. Fuchs, I. Iuliani, V. Firmano, G. Tallarico, L. Calabrese, J. Grilli, B. Sclavi, L. Ciandrini, and M. C. Lagomarsino, Incoherent feedback from coupled amino acids and ribosome pools generates damped oscillations in growing E. coli, Nature Communications **16**, 3063 (2025).

[6] D. W. Erickson, S. J. Schink, V. Patsalo, J. R. Williamson, U. Gerland, and T. Hwa, A global resource allocation strategy governs growth transition kinetics of Escherichia coli, Nature 10.1038/nature24299 (2017).

[7] G. Chure and J. Cremer, An optimal regulation of fluxes dictates microbial growth in and out of steady state, eLife **12**, e84878 (2023).

[8] J. C. Kratz and S. Banerjee, Dynamic proteome trade-offs regulate bacterial cell size and growth in fluctuating nutrient environments, Communications Biology **6**, 486 (2023).

[9] D. Goberman, A. Roy, and R. Pugatch, Slowing translation to avoid cellular depletion of active ribosomes at slow growth rates, Physical Review Research **8**, 013226 (2026), 2503.07837.

[10] G.-W. Li, D. Burkhardt, C. Gross, and J. Weissman, Quantifying Absolute Protein Synthesis Rates Reveals Principles Underlying Allocation of Cellular Resources, Cell **157**, 624 (2014).

[11] A. Schmidt, K. Kochanowski, S. Vedelaar, E. Ahrné, B. Volkmer, L. Callipo, K. Knoops, M. Bauer, R. Aebersold, and M. Heinemann, The quantitative and condition-dependent Es-

cherichia coli proteome, Nature Biotechnology **34**, 104 (2016), #proteome #proteome #proteome #proteome.
